# MELODY: Mediation Analysis in Logistic Regression for High-Dimensional Mediators and a Binary Outcome

**DOI:** 10.1101/2025.11.12.688038

**Authors:** Sunyi Chi, Xingyu Li, Peng Wei, Xuelin Huang

**Affiliations:** Department of Biostatistics, The University of Texas MD Anderson Cancer Center, Houston, TX, USA

**Keywords:** binary outcome, high-dimensional mediators, mediation analysis, metabolomics, total mediation effect, transcriptomics

## Abstract

Mediation analysis is a pivotal tool for elucidating the indirect effect of an environmental factor or treatment on disease through potentially high-dimensional omics data, such as gene expression profiles. However, traditional mediation analysis methods tailored for binary outcomes often rely on the rare disease assumption in logistic regression and provide inadequate measures of total mediation effect when multiple mediators have effects in different directions. In this paper, we develop a MEdiation analysis framework in LOgistic regression for high-Dimensional mediators and a binarY outcome (MELODY). It leverages a second-moment-based measure analogous to the ***R***^***2***^ for linear models to quantify the total mediation effect. We also develop a variable selection procedure for high-dimensional data to reduce bias introduced by non-mediators. Our comprehensive simulations demonstrate the superior performance of MELODY in scenarios with non-rare disease binary outcomes and high-dimensional mediators. We apply MELODY to the Framingham Heart Study of over 5000 individuals to analyze the mediation effects of metabolomics and transcriptomics data on the pathways from sex to incident coronary heart disease.

## 1 Introduction

Mediation analysis is a commonly used statistical approach that aims to determine the indirect effect of an environmental factor or phenotype variable on diseases of interest through potential intermediate variables (Cashin et al., 2023). Typically, the targeted disease variable (i.e., dependent variable) is a binary variable related to disease diagnosis. The effect of multiple mediators and high-dimensional omics mediators has received much attention (Clark-Boucher et al., 2023; Fang et al., 2021; Sohn and Li, 2019; Zhang et al., 2018; Zeng et al., 2021). In a causal framework, formal definitions of total, direct, and indirect effects were first introduced by Robins et al. (1992) and Pearl (2001). For binary outcomes, VanderWeele and Vansteelandt (2010) and Valeri and VanderWeele (2013) derived product-based measures on the odds ratio scale for a rare binary outcome with a prevalence of below approximately 5%. This requires the assumption that the product measure on the odds ratio scale can be approximated by the difference measure on the relative risk scale in estimating the mediation effects. Existing mediation analysis methods based on these traditional effect measures are limited in handling high-dimensional mediation analyses with binary outcomes (Rijnhart et al., 2019). First, the rare outcome assumption may not hold in practice since some diseases, e.g., cardiovascular disease, are common with high prevalence and incidence rates. Second, the product-based effect measures are on additive scale, leading to mediators’ effects of different directions being cancelled out in the multiple-mediator model.

Several approaches for mediation analysis with a common dichotomous outcome have been proposed without assuming the rarity of the binary outcome. Doretti et al. (2022) and Samoilenko et al. (2021) have derived exact parametric measures on the odds ratio scale that do not require the rare outcome assumption for settings with a binary outcome and a binary mediator. Other studies developed probit models (Gaynor et al., 2019; Imai et al., 2010; Sohn et al., 2022) or a student-t (t-link) approximation under a Bayesian framework (Caubet et al., 2023) to relax the rare disease assumption. However, these methods are developed in single-mediator models for binary outcomes and cannot estimate the total mediation effect accommodating multiple or high-dimensional mediators. Loh et al. (2022), Liu et al. (2022), and Derkach et al. (2019) proposed methods for high-dimensional mediation analysis but still have the rare outcome assumption for binary outcomes. Nguyen et al. (2016) and Lai et al. (2020) developed methods in the probit model for multiple mediators on a binary outcome and do not require the rare outcome assumption. However, these methods still used the risk-difference- and odds-ratio-based effect measures for multiple or high-dimensional mediators, which could cancel out conflicting individual mediators’ effects in the estimation of the total mediation effect.

Here we address these limitations by exploring *R*^2^-based high-dimensional mediation analysis for a binary outcome. As a complementary alternative to the traditional mediation effect measures based on the mean (first moment), MacKinnon (2008) and Fairchild et al. (2009) proposed the *R*^2^-based mediation effect size measure in the linear model. Recently, Yang et al. (2021), Shi et al. (2022), Xu et al. (2024), and Chi et al. (2024) extended the use of the *R*^2^ measure to multiple- or high-dimensional mediator models for continuous outcomes and survival outcomes. The *R*^2^-based effect measure can capture the nonzero total mediation effect in the presence of bi-directional component-wise mediation effects, likely ubiquitous in high-dimensional omics data settings (Xue et al. (2022)). Moreover, a mediation analysis framework based on a *R*^2^-based effect measure does not require the rare outcome assumption. However, the *R*^2^-based mediation analysis method for high-dimensional mediators and binary outcomes has yet to be studied, and there is no consensus on the definition of *R*^2^ in logistic regression.

To fill the knowledge gap described above, we first evaluated and compared various *R*^2^ measures in the logistic regression framework for mediation analysis. Once having identified the most suitable *R*^2^-based effect size measure, we proposed a MEdiation analysis framework in LOgistic regression for high-Dimensional mediators and a binarY outcome (MELODY). To evaluate the bias, variance, and mediator selection accuracy of our mediation effect estimation, we conducted extensive simulation studies in various settings by varying the prevalence of disease, the number of putative mediators, the effect sizes, the types of non-mediators, and the dimension of data. We demonstrated the superior performance of MELODY in scenarios with non-rare disease binary outcomes and high-dimensional mediators. Furthermore, we applied MELODY to the Framingham Heart Study to analyze the mediation effects of transcriptomics and metabolomics data on the pathways from sex to incident coronary heart disease.

## 2 Methods

### 2.1 Mediation Models

Let *X* be the exposure, *M* be a set of *p* mediators {*M*_1_, *M*_2_, …, *M*_*p*_}, and *Y* be a binary outcome, as illustrated in Figure 1. Let *Z* denote baseline covariates such as age, which may be a potential confounder. To obtain a closed-form expression for the mediation effects, we use logistic regression to model the binary outcome. The mediation model is therefore defined using the following four logistic regression models:

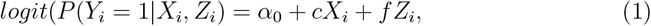

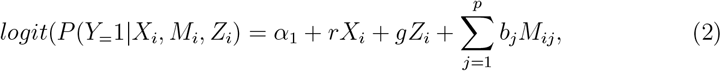

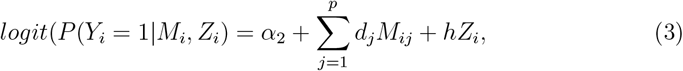

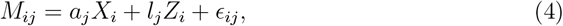

where *logit*(*x*) = *log*(*x/*(1 − *x*)) for *x* ∈ (0, 1) is the logit function. *M*_*i*_ = (*M*_*i*1_, *M*_*i*2_, …, *M*_*ip*_)^*′*^ is the *p*-dimensional mediator vector for subject *i* = 1, 2, 3, …, *n* and, without loss of generality, *X, M*_*j*_ and *Z* are standardized to have mean 0 and variance 1. Equations (1), (2), and (3) are logistic regression models describing the relationship (1) between *Y* and *X* given *Z*, (2) between *Y, X*, and *M* given *Z*, and (3) between *Y* and *M* given *Z*, respectively. *c* is the parameter relating the exposure to the outcome; *r* is the direct effect parameter relating the exposure to the outcome; *b*, defined as (*b*_1_, …, *b*_*p*_)^*′*^, is the parameter vector relating the mediators to the outcome with adjustment for the effect of the exposure; and *d* = (*d*_1_, …, *d*_*p*_)^*′*^, is the parameter vector relating the mediators to the outcome. equation (4) characterizes how exposure influences the mediators, where *a* = (*a*_1_, …, *a*_*p*_)^*′*^ is the parameter vector relating the exposure to the mediators, *l* ∈ *R*^*p*∗*γ*^, *g* ∈ *R*^*γ*^ (*γ* is number of covariates *Z*), and residual *ϵ*_*i*_ = (*ϵ*_*i*1_, *ϵ*_*i*2_, …, *ϵ*_*ip*_)^*′*^ ∼ *MV N* (0, *E*_*p×p*_), for *i* = 1, …, *n*.

**Fig. 1.**
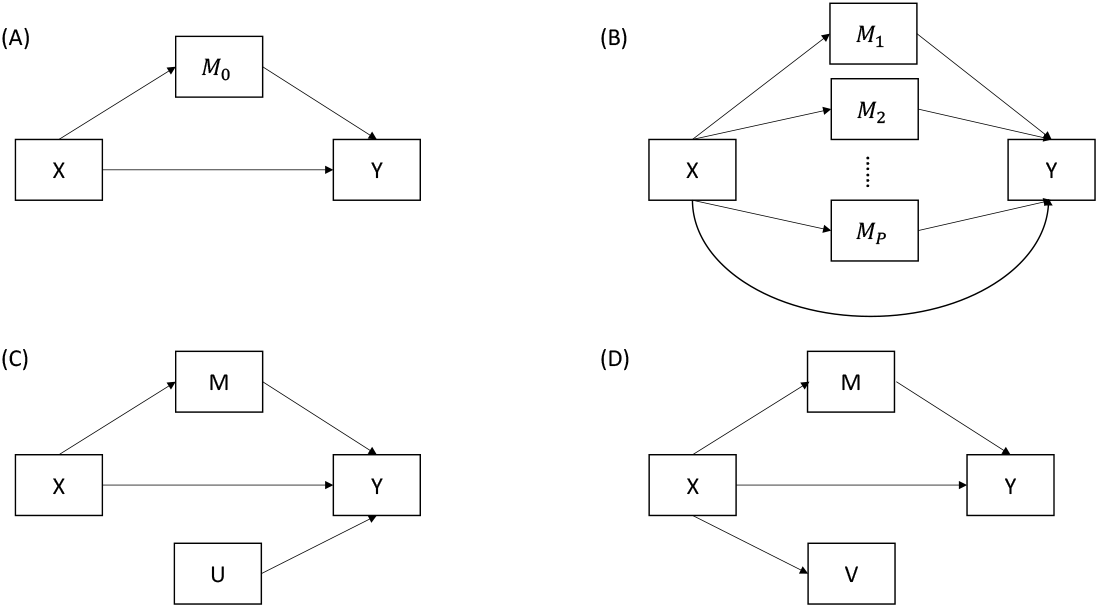
Diagrams for potential mediation settings. *X* is the independent variable, *Y* is the binary outcome, and *M* is a set of true mediator. *U* is a set of variables associated with *Y* but not *X*, and *V* is a set of variables associated with *X* but not *Y* . (A) A single-mediator model. (B) A multiple-mediator model. (C) A model with non-mediator *U* . (D) A model with non-mediator *V* .

To formally define the causal effect in the counterfactual framework (Pearl, 2001; Robins et al., 1992), we will specifically follow the notations of VanderWeele and Vansteelandt (2010). Let *Y*_*x*_ and *M*_*x*_ be an individual’s outcome *Y* and mediators *M*, respectively, that would have occurred had the exposure been *X* = *x*. Further, denote *Y*_*xm*_ as an individual’s outcome *Y* under exposure *X* = *x* and mediators *M* = *m*. The natural direct effect (NDE) on the odds ratio scale has been previously defined. It compares the odds of the outcome with exposure *x* and mediator *M*_*x*_∗ to the odds of the outcome with exposure *x*^∗^ and mediator still *M*_*x*_∗, conditional on any covariates *Z* = *z*. On the odds ratio scale, the natural indirect effect (NIE) is a measure of the effect on the outcome through the mediator and, thus, compares the odds of the outcome with exposure *x* and mediator *M*_*x*_ to the odds of the outcome with exposure *x* and mediator *M*_*x*_∗ . If assumptions from VanderWeele and Vansteelandt (2010) hold and regression models (2) and (4) are correctly specified, then the NDE and NIE odds ratios are given by

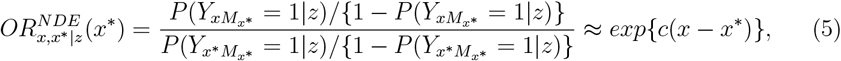

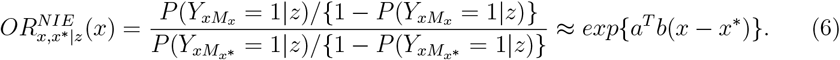

These expressions, referred to as the “Baron-Kenny” approach in a single-mediator model (Baron et al., 1986) and the product measure in a multiple-mediators model, essentially use *c* for the direct effect and 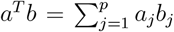 for the indirect effect. A related common approach, referred to as the difference measure, consists of regression models (1) and (2), examining the difference between coefficients for *X, c* − *r*, as the indirect effect. The traditional proportion method then uses (*c* − *r*)*/c* as the measure of indirect effect.

The difference and product measures coincide for continuous outcomes but not dichotomous outcomes (Fritz et al., 2008; Wang, 2018). VanderWeele and Vansteelandt (2010) showed that only with a rare outcome are these two approximately equal and able to provide a formal counterfactual interpretation for dichotomous outcome. Furthermore, in high-dimensional mediation analysis, the component-wise mediation effects (*a*_*j*_*b*_*j*_’s) could cancel out due to the likely presence of opposite mediation directions, resulting in close to zero or negative product measures. However, existing mediation analysis methods for multiple mediators and dichotomous outcomes still use the product measure. Therefore, our study focuses on developing a mediation analysis framework for binary outcomes and high-dimensional mediators that does not require the rare disease assumption and does not cancel the component-wise mediation effects *a*_*j*_*b*_*j*_’s in different directions in the total mediation effect estimation.

### 2.2 Proposed MELODY for high-dimensional mediation analysis with a binary outcome

In this section, we propose a high-dimensional mediation analysis method, MELODY, based on a second-moment indirect effect measure for dichotomous outcome inspired by Fairchild et al. (2009). To introduce the details of the proposed method MELODY, we first discuss the proposed *R*^2^-based mediation effect measure, and then introduce the mediator variable selection and the complete workflow in the following section.

#### 2.2.1 Proposed *R*^2^ measures for multiple-mediator mediation analysis under the logistic regression

The *R*^2^ mediation effect measure, which was proposed in a single-mediator model for a continuous outcome by Fairchild et al. (2009), has good performance with high-dimensional and complex structured data (Xu et al., 2024; Xu and Wei, 2024; Yang et al., 2021) and survival outcomes with modified *R*^2^ for time-to-event data (Chi et al., 2024; Shi et al., 2022). Let *MED* be the measure of mediation effect, which is defined by the amount of variation in outcome *Y* that is explained by exposure *X* through mediators *M* as follows:

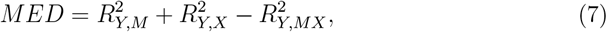

where 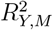 is the proportion of variation in outcome *Y* explained by the mediators, 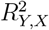 is the variation in *Y* explained by the exposure, and 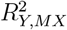 is the proportion of variation in *Y* explained by both the mediators and the exposure. Thus, 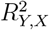 quantifies the total effect, whereas 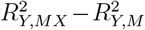 is the direct effect. Therefore, the *R*^2^ measure of indirect effect *MED* is the total effect minus the direct effect, which is given in equation (7). The shared over simple effect (SOS) defined as 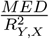 by Lindenberger et al. (1998) is the standardized exposure-related variation in the outcome that is shared with the mediators.

Motivated by the *R*^2^-based mediation effect measure for continuous and time-to-event outcomes, next we formally extend *R*^2^-like mediation effect measures to mediation models with binary outcomes. There are many ways to calculate a pseudo-*R*^2^ for logistic regression, with no consensus on which one is the best. In the context of mediation analysis, particularly for logistic regression models, selecting appropriate *R*^2^ measures is crucial. Our focus narrows to three prominent *R*^2^ measures: McFadden’s *R*^2^ (McFadden, 1973), Nagelkerke’s *R*^2^ (Nagelkerke et al., 1991), and Tjur’s *R*^2^ (Tjur et al., 2009). These measures were identified based on comprehensive review, theoretical and numerical evaluations in various studies, fulfilling essential criteria for effective *R*^2^ measures in logistic regression (Kvalseth et al., 1985; Royston et al., 2006; Schemper et al., 1996).

McFadden’s *R*^2^ was highlighted for its robust performance in multiple logistic regression (Menard et al., 2000) and has been deemed attractive in the field (Allison, 2014). Specifically, it is defined as

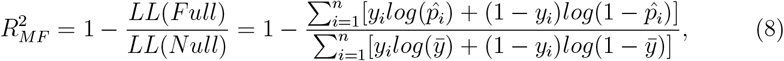

where *LL*(*Full*) is the log-likelihood function of a specified model of interest, *LL*(*Null*) is the log-likelihood of an intercept-only model, and 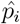 is predicted by the model of interest, and 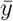 is the mean of all *y*_*i*_’s. Plugging 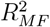 into equation (7) results in our proposed mediation effect measure *MED* for high-dimensional mediation analysis for binary outcomes; see Supplementary Materials Section 1 for details. Nagelkerke’s *R*^2^ is known for its compatibility and close agreement with the *R*^2^ of the general linear model, offering an intuitively clear interpretation (Schemper et al., 1996). Both McFadden and Tjur’s measures were found useful in epidemiological models assessing disease risk (Hughes et al., 2019). Our study considers these three *R*^2^ measures in the mediation analysis framework to evaluate their applicability and robustness in high-dimensional mediation analysis.

The comparative analysis of these three *R*^2^ measures, based on our extensive simulation studies, is detailed in the Supplemental Materials Section 2. Briefly, we evaluated the the performance of McFadden’s *R*^2^, Nagelkerke’s *R*^2^, Tjur’s *R*^2^, the product, and the difference measures with respect to the disease prevalence, mediation effect size and the number of mediators. Firstly, as shown in Supplemental Figure 1, McFadden’s *R*^2^-based mediation measure exhibited stability and independence from disease prevalence, ranging from rare to common, compared with Nagelkerke’s *R*^2^, Tjur’s *R*^2^ and the difference measure, which is an appealing property for binary outcomes (Royston et al., 2006). Secondly, McFadden’s *R*^2^ measure consistently showed lower values across the board, indicating a more conservative estimate of variance explained, which aligns with its known properties in the literature (Allison, 2014) (Supplemental Figures 1-3). Thirdly, as effected, all mediation effect measures under comparison increased with increasing mediation effect (parameter *b* in equation (2)) and increasing number of mediators (Supplemental Figures 2 and 3, respectively). Finally, the difference between the product measure and the difference measure increased as the prevalence of disease increased up to 50%, which aligns with its rare disease assumption (Supplemental Figure 1). Based on the simulation study, we therefore advocate for the use of McFadden’s *R*^2^ in logistic models for more robust and stable mediation effect estimation.

### 2.3 Partial *R*^2^ to control confounding effects

As illustrated by VanderWeele et al. (2016), failure to control for exposure-outcome, exposure-mediator, and mediator-outcome confounding effects in mediation analysis can substantially bias estimates of mediation effects, whereas adjusting for measured confounders as covariates in mediation models can reduce the bias. Therefore, we propose using the partial *R*^2^ measure motivated by Chi et al. (2024) to adjust for potential confounders in mediation analysis for binary outcomes. In general, partial *R*^2^ is defined as the proportion of variation in the outcome variable that cannot be explained in a reduced model but can be explained by the predictors specified in a complete model. To adjust for the covariates in the indirect effect of exposure on the outcome through mediators, partial *R*^2^ is adapted based on our original formulas to calculate the mediation effect as shown below.

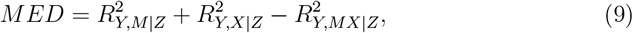

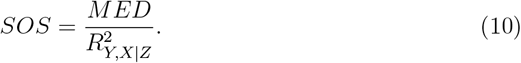

According to the definition of partial *R*^2^, 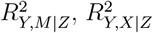, and 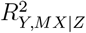 are

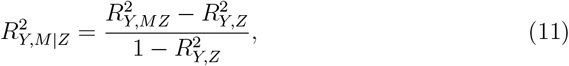

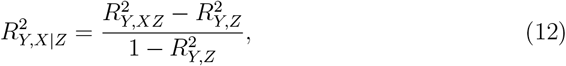

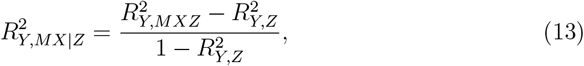

where 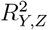 is McFadden’s *R*^2^ measure for *logit*(*P* (*Y*_*i*_ = 1|*Z*_*i*_) = *α*_3_ + *kZ*_*i*_. Therefore, 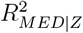 can be interpreted as the variation of the outcome variable that the exposure variable can explain through mediators adjusting for confounder(s) *Z*.

#### 2.3.1 Mediator Variable Selection

Although we propose a method to control for confounding variables by partial *R*^2^, the following assumptions are made as in VanderWeele et al. (2014): conditional on covariates *Z*, (1) no unmeasured confounders between the exposure and the outcome, between the mediators and the outcome or between the exposure and the mediators and (2) no exposure-induced confounding between the mediators and the outcome.

To perform mediation analysis of high-dimensional mediators, we must first select the true mediators in the pathway from the exposure to the outcome before estimating the mediation effect. Let **S**_**0**_ denote a set of potential mediators, **M** denote a set of true mediators, **U** denote a set of variables associated with the outcome but not the exposure (Fig. 1C), and **V** denote a set of variables associated with the exposure but not the outcome (Fig. 1D). Inclusion of the **U, V** or noise variables in mediation models may bias the effect size estimation as to be illustrated later in the simulation study.

Mediator selection is a new challenge for high-dimensional mediation analysis. In the traditional mediation analysis, the putative mediating variables are selected based on *a priori* subject-matter knowledge or hypothesis testing (Cashin et al., 2023). However, the true mediators are hardly known *a priori* and usually sparse in the high-dimensional mediation analysis setting (Chi et al., 2024; Yang et al., 2021). As shown in Fig. 2, we randomly split the whole dataset into 75%:25%: the first part for mediator variable selection and the second part for mediation effect estimation based on the selected mediators to overcome the “winner’s curse” in high-dimensional variable selection (Zhong and Prentice, 2008). Specifically, for variable selection we adopt the sure independence screening (Fan et al., 2008) with the minimax concave penalty (MCP) (Zhang, 2010) to reduce dimension and exclude V-type non-mediators and use and the false discovery rate (FDR) adjusted p-value based on the Benjamini-Hochberg procedure (Benjamini et al., 1995) to filter out U-type non-mediators as inspired by Chi et al. (2024). We examined each step’s effects and tested the FDR thresholds in the simulation studies.

**Fig. 2.**
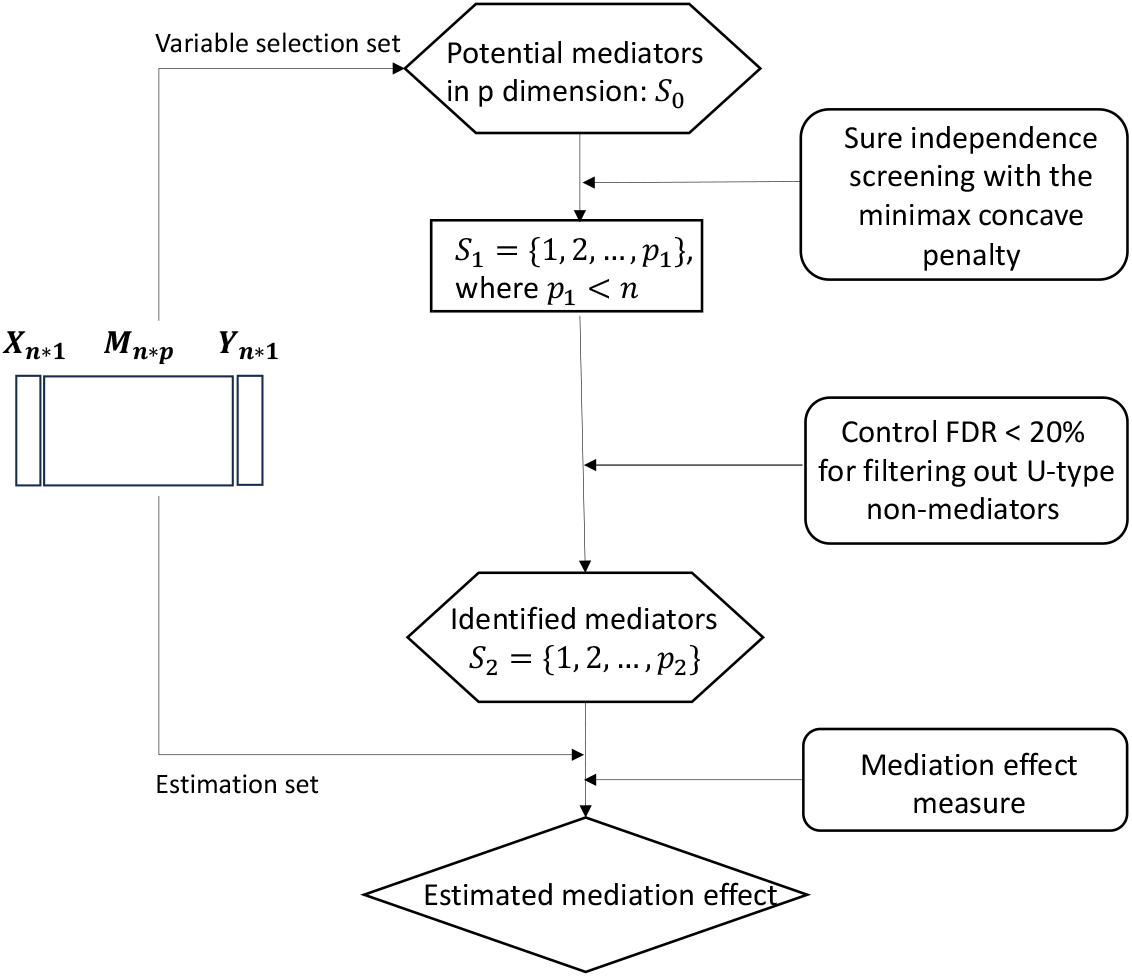
Overall workflow of MELODY. The workflow consists of four main steps: (a) sure independence screening for preliminary screening, (b) minimax concave penalty for variable selection, (c) false discovery rate (FDR) control at 0.2 level for variable selection, and (d) mediation effect estimation using *R*^2^-based measures.

## 3 Simulations

We investigated the performance of the proposed method MELODY compared with existing mediation effect size measures for logistic regression and assessed the mediator selection methods for high-dimensional mediators with binary outcomes in the logistic regression framework.

### 3.1 Simulation Design

We simulated the data following the models in equations (2) and (4). Specifically, the exposure X, confounding variables Z, noise mediators, U-type non-mediators, and error term *ϵ*_*ij*_ were randomly generated from the standard normal distribution N(0, 1) for subject *i* = 1, …, *n*. Then, the true mediators and V-type non-mediators were generated using equation (4), and the binary outcome was generated from equation (2).

To evaluate the performance of the proposed mediation effect measure, we calculated the bias and variance of mediation effect estimators. The bias of an estimator was calculated by 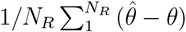, where 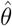 was the estimator of the mediation effect measure *θ* and *N*_*R*_ was the number of replicates in simulation. For a fair comparison, we evaluated different effect measures by relative bias instead of bias. The relative bias is calculated by 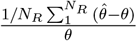 . Our results are summarized based on *N*_*R*_ = 500 replications. We calculated a pseudo-true mediation effect with known parameters and true mediators in our simulation data-generation model. Note that the pseudo-true *R*^2^ mediation effect is not the exact true value because the true parameters c and d are unknown. As a result, the *R*^2^ of models (2) and (4) (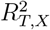 and 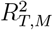) are estimated values based on a very large sample size (*n* = 200, 000) to circumvent the undue influence of model fitting uncertainty following (Chi et al., 2024).

The existence of either U or V-type non-mediators could bias the estimation of mediation effects. Thus, we evaluated variable selection methods: A) filter out V-type non-mediators by sure independent screening (SIS) with MCP on model (2); B) filter out V-type non-mediators by SIS with MCP on model (2) and filter out U-type non-mediators by controlling FDR *<*0.2; C) filter out V-type non-mediators by SIS with MCP on model (2) and filter out U-type non-mediators by controling FDR *<*0.05; D) not filter out any non-mediator. We evaluated the mediator selection performance by examining their impact on the bias and standard deviation (SD) of the mediation effect estimations and via sensitivity and specificity as: 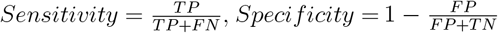. Specifically, the sensitivity, also named true positive rate (TPR), refers to the method’s ability to detect mediators out of all true mediators, and specificity, which is 1 - false positive rate (FPR), represents the proportion of non-mediators selected from all true non-mediators. A method with a higher sensitivity and specificity selects mediators with a higher accuracy and leads to less bias in mediation effect estimation. Under the high-dimensional mediators and binary outcome framework, we conducted simulation studies in the following seven settings:

#### Setting I: Varying effect sizes (*ab*)

We set the sample size *n* = 2000, number of true mediators *m* = 5, number of potential mediators *p* = 1000, *r* = 1, and *α*_1_ = 0; we varied the value of coefficients *a* and *b* in setting I, such that *ab* = 0.5, 0.4, 0.3, 0.2, 0.1, respectively.

#### Setting II: Varying prevalence of the disease (pr)

We set the sample size *n* = 2000, number of true mediators *m* = 5, number of potential mediators *p* = 1000, and *r* = 1. We set *a* = 0.4 and *b* = 0.25 for each true mediator. We varied the value of coefficient 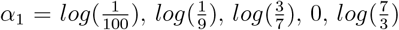, *log*(9) such that *pr* = 0.029, 0.052, 0.175, 0.276, 0.358, respectively.

#### Setting III: Presence of U- or V-type non-mediators

We set the sample size *n* = 2000, number of true mediators *m* = 5, number of potential mediators *p* = 1000, *r* = 1, and *α*_1_ = 0. We set *a* = 0.2 and *b* = 0.3 for each true mediator. In addition to noise variables, we introduced 10 U- or V-types of non-mediators for each setting. For U-type non-mediators, we set *a* = 0 and *b* = 0.3. For V-type non-mediators, we set *a* = 0.2 and *b* = 0.

#### Setting IV: Varying number of true mediators

We set the sample size *n* = 2000, number of potential mediators *p* = 1000, *r* = 1, and *α*_1_ = 0. We set *a* = 0.4 and *b* = 0.25 for each true mediator. We varied the number of true mediators *m* = 5, 20, 50.

#### Setting V: Varying dimension of the data (n/p)

We set the sample size *n* = 2000, number of true mediators *m* = 5, *r* = 1, and *α*_1_ = 0. We set *a* = 0.4 and *b* = 0.25 for each true mediator. We varied the number of potential mediators *p* = 1000, 2000, 5000, and 10000.

#### Setting VI Varying confounding effects

We set the sample size *n* = 2000, number of true mediators *m* = 5, number of potential mediators *p* = 1000, *r* = 1, and *α*_1_ = 0. We set *a* = 0.2 and *b* = 0.3 for each true mediator. For covariates *Z*, we set the parameters *l* = 0.1, and *g* = 0.1, 0.5, 0.75, and 1.

#### Setting VII: Conflicting directions of component-wise mediation effects

We set the sample size *n* = 2000, number of true mediators *m* = 5, number of potential mediators *p* = 1000, *r* = 1, *α*_1_ = 0, we set *a* = 0.4 for each true mediator. We varied the directions of the component-wise mediation effects to study their impact on the total mediation effects by different measures. In the same direction setting, we set *b* = 0.25 for the five true mediators such that all true mediators had positive effects. In the conflicting directions setting, we set *b* = (0.5, 0.5, 0.5, -0.25, -0.25) such that three mediators had positive effects while the other two had negative ones.

### 3.2 Simulation Results

Fig. 3-5 present the relative bias and variance under the high-dimensional settings I to VII. The proposed MELODY using *MED* as the mediation effect measure had a smaller variance in all settings compared to the product measure and difference measure as shown in Fig. 3-5. Figure 3A shows that the difference measure had higher bias as the effect sizes decreased, while the MELODY and product measures were relatively stable with varying effect sizes. Figure 3B, along with Supplemental Fig 3, shows that the difference measure and product measure were approximately equal only under a very low prevalence of disease. When the prevalence of disease was larger than 0.175, these two measures were substantially different. Figure 4A shows that the proposed *MED* estimator had minimal bias with V-type non-mediators but were slightly biased and had smaller variance than did the product and difference measures with U-type non-mediators. Figure 4B shows that MELODY had stable performance in setting VII when individual mediators had conflicting directions of effects. However, the product and difference measures introduced more bias under this setting. We further confirmed that the proposed *MED* was more robust in terms of bias and variance under high-dimensional settings with varying numbers of true mediators and potential mediators, where the performance of the difference measure deteriorated under these challenging settings (Fig. 4C and 5A). In addition, MELODY and the product measure had more negligible bias than did the difference measure in the presence confounding variables and varying confounding effects (Fig. 5B).

**Fig. 3.**
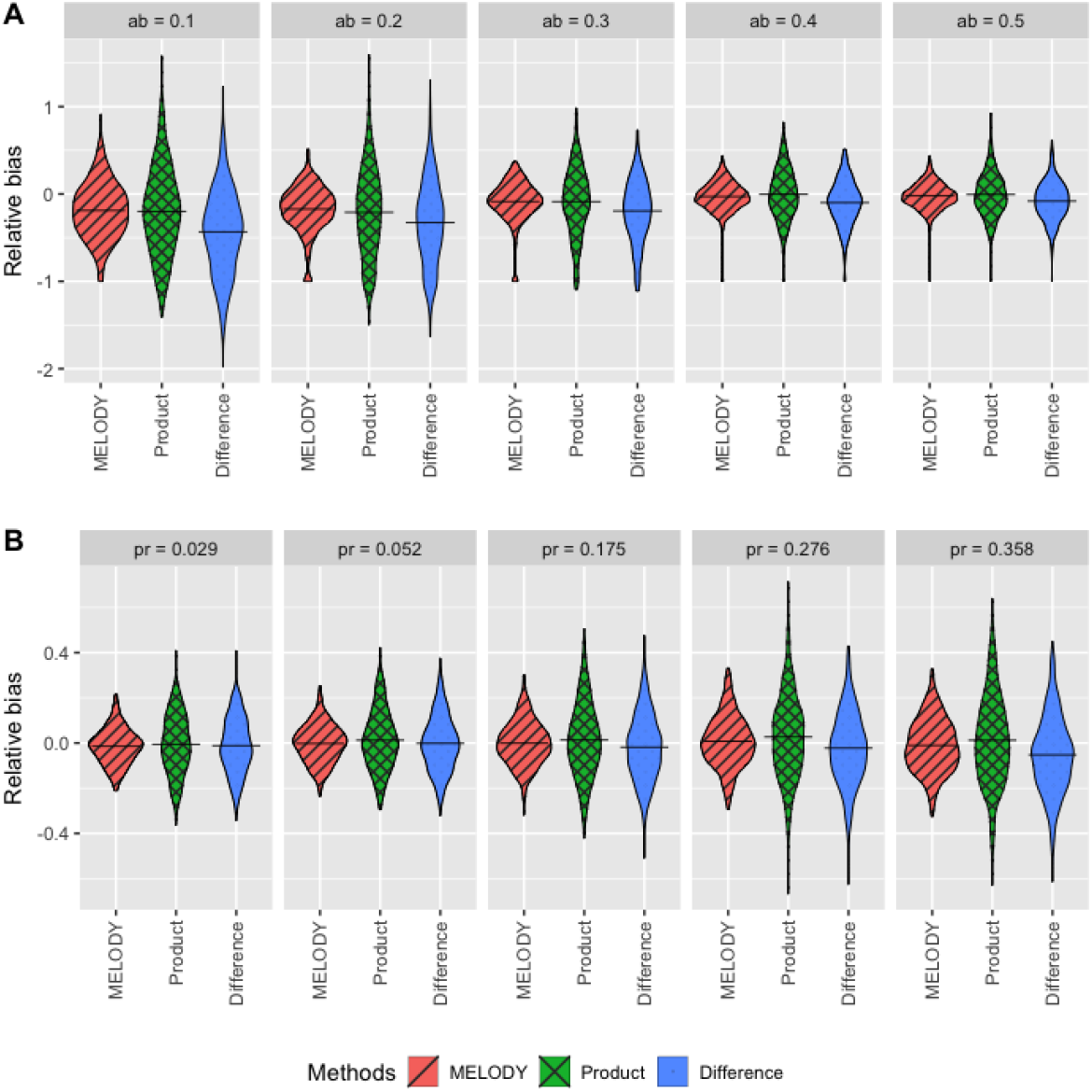
Violin plots of the relative bias of MELODY, product, and difference methods under simulation settings I and II. A Setting I: Varying effect sizes *ab* across 0.5, 0.4, 0.3, 0.2, and 0.1. B Setting II: Varying revalence of the disease *pr* across 0.029, 0.052, 0.175, 0.276, and 0.358. The x-axis corresponds to three methods: the proposed MELODY, the product method, and the difference method. The y-axis corresponds to the relative distance between estimated values and true values as 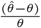 . The value of the crossbar is the mean across 500 simulation replications, representing the relative bias.

**Fig. 4.**
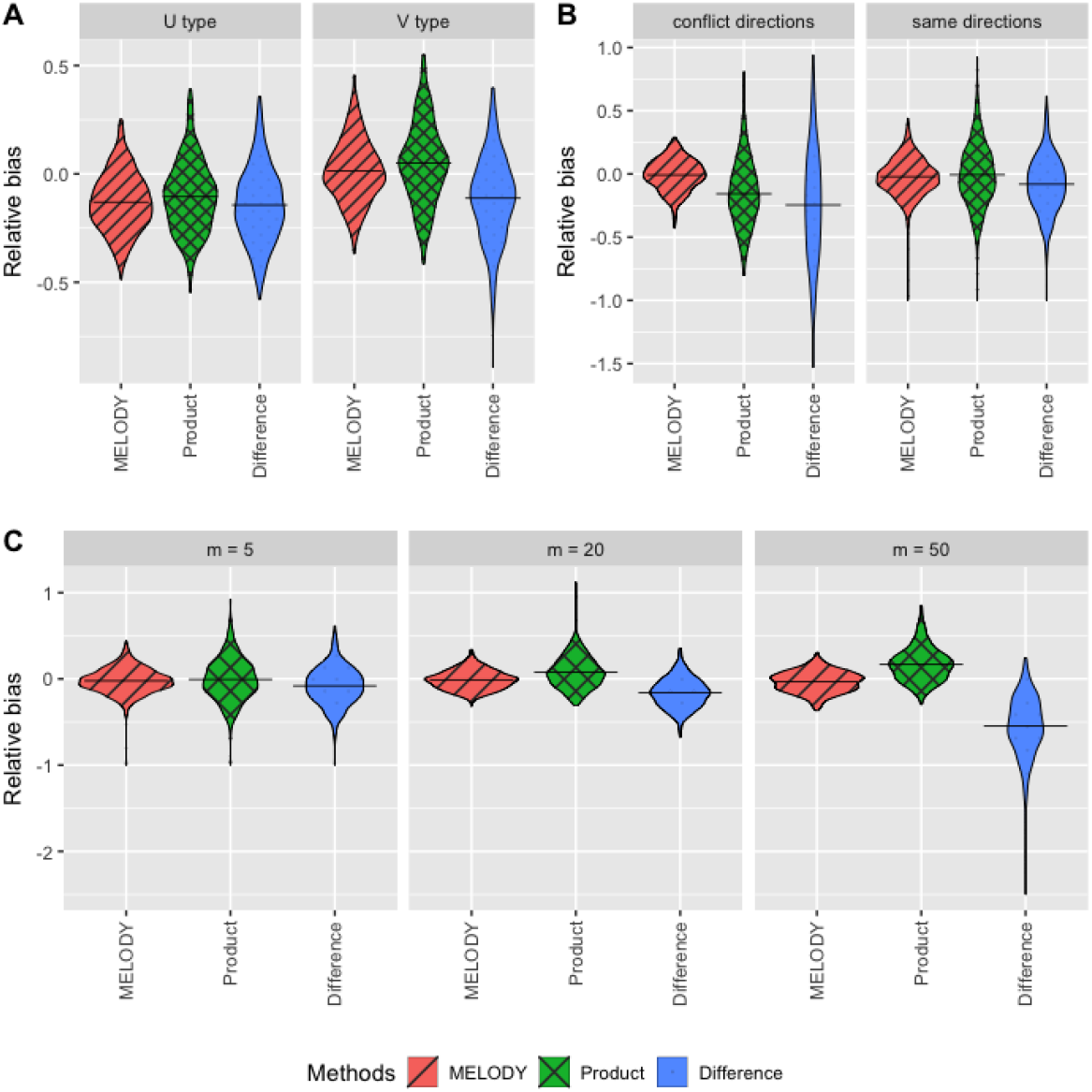
Violin plots of the relative bias of MELODY, product, and difference methods under simulation settings III, IV, and VII. A Setting III: U- or V-type non-mediators exist in addition to noise variables. For U-type non-mediators, we set *a* = 0, *b* = 0.3. For V-type non-mediators, we set *a* = 0.2, *b* = 0. B Setting VII: In the same direction setting, all five true mediators had positive component-wise mediation effects. In the conflicting directions setting, we changed the effect directions of 2 true mediators from positive to negative. C Setting IV: Varying number of true mediators *m* across 5, 20, and 50. The x-axis corresponds to three methods: the proposed MELODY, the product method, and the difference method. The y-axis corresponds to the relative distance between estimated values and true values are 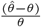 . The value of the crossbar is the mean across 500 simulation replications, representing the relative bias.

**Fig. 5.**
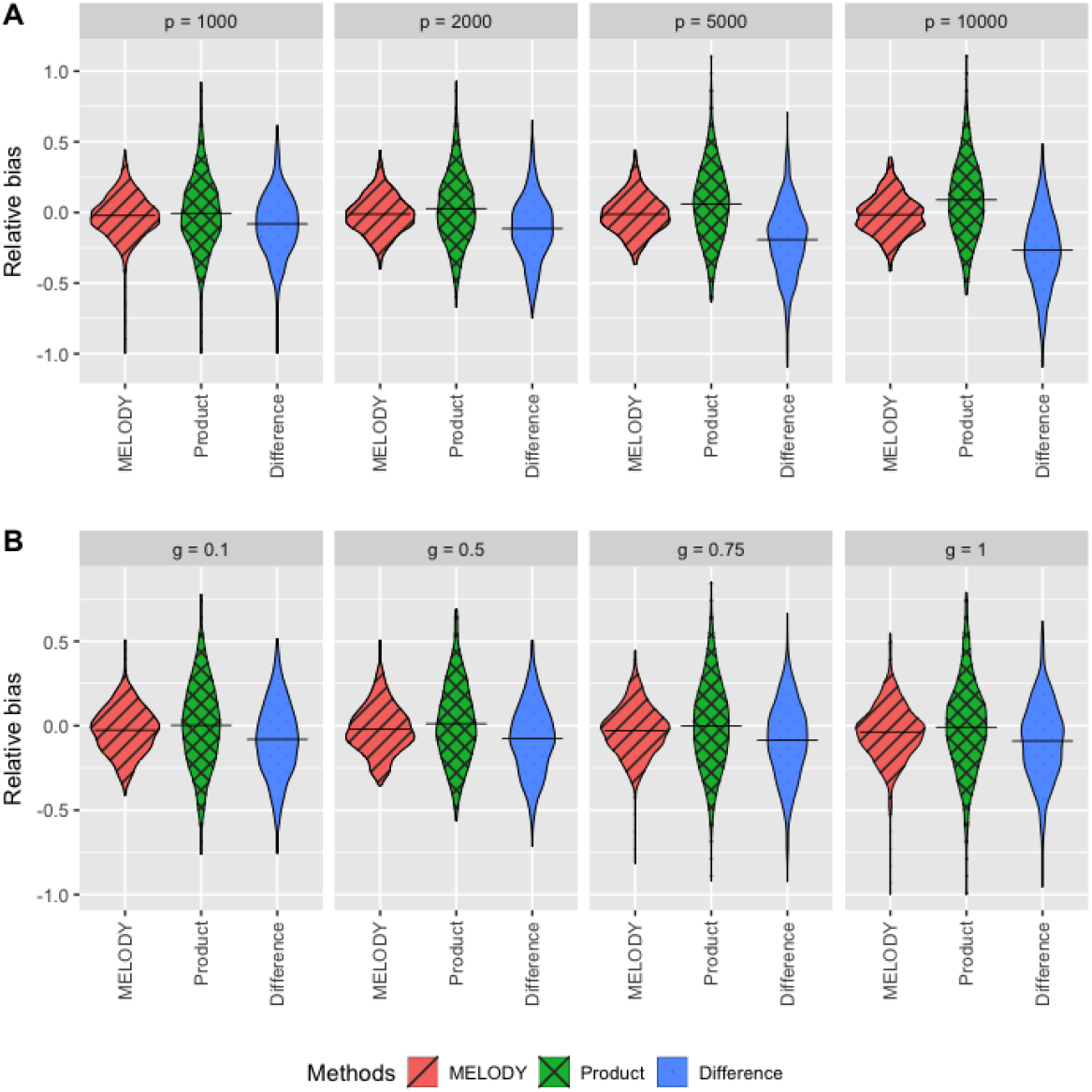
Violin plots of the relative bias of MELODY, product, and difference methods under simulation settings V and VI. A Setting V: The number of potential mediators p are varied across 1000, 2000, 5000, and 10000. B Setting VI: Confounding variable exists and the confounding effects are varied by varying the parameter *g* across 0.1, 0.5, 0.75, and 1. The x-axis corresponds to three methods: the proposed MELODY, the product method, and the difference method. The y-axis corresponds to the relative distance between estimated values and true values 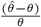. The value of the crossbar is the mean across 500 simulation replications, representing the relative bias.

In addition, we present the results of the seven simulation settings in detail in Supplemental Tables S1 - S7. Settings in Table S1, S2, and S4-S6 represent the scenarios of high-dimensional settings with fixed sample size *n* = 2000 and number of potential mediators *p* = 1000, but we varied the effect sizes of mediators, the prevalence of disease, the number of true mediators, the dimensions of data, and the confounding effect sizes, respectively. Table S3 shows the results of setting III, where we introduced 10 U- or V-type non-mediators for each setting besides the noise variables to evaluate the effect of non-mediators U and V. In setting VII, presented in Table S7, we let the true mediators have different directions of mediation effects. Upon comparison of the mediator selection methods shown in Tables S1 - S7, the mediator selection procedure with or without FDR barely affected the bias of mediation effect estimator 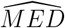. However, FDR with a more conservative threshold increased the specificity. Considering the interpretation and follow-up analysis of the selected mediators, we will use SIS with MCP to filter out V-type non-mediators and control FDR *<*0.2 to filter out U-type non-mediators.

## 4 Application to the Framingham Heart Study

To further evaluate the proposed MELODY method, we applied it to the Framingham Heart Study (FHS) of over 4000 individuals to analyze the mediation effects of transcriptomics (gene expression) and metabolomics on the pathways from sex to incident coronary heart disease.

Coronary heart disease (CHD) is a leading cause of death worldwide, with men being at a higher risk of CHD than women (Tsao et al., 2022). Studies have found that the expression of candidate genes related to coronary heart disease is also associated with sex. We were interested in exploring how metabolomics and gene expression (GE) mediate the effect of sex on CHD risk in the FHS, which is a prospective cohort study of cardiovascular disease that has been ongoing since the 1940s and includes three generations: the Original Cohort, the Offspring Cohort, and the Third Generation Cohort (Splansky et al., 2007). We downloaded exposure, metabolomics, and outcome data of 1902 Offspring Cohort individuals from NIH/NCBI dbGaP, 317 of whom were diagnosed with CHD (prevalence of CHD: 16%) during follow-up after Exam 5 when plasma was collected for metabolomics profiling. A total of 190 metabolites were profiled as potential mediators in our analysis. We also downloaded the genome-wide gene expression, i.e., the transcriptome data of 5204 participants in the Offspring and 3rd Generation Cohorts. Gene expression of 17873 genes were profiled using the Affymetrix GeneChip Human Exon 1.0 ST Array platform based on blood sample collected at Exam 8 for the Offspring Cohort and Exam 2 for the 3rd Generation Cohort. In total 181 participants were diagnosed with CHD during follow-up after Exam 8 (Offspring Cohort) and Exam 2 (the 3rd Generation Cohort) (3.48% prevalence). The exposure variable was sex. The binary outcome was whether CHD was diagnosed during the follow up, ascertained via clinical exams or annual telephone surveys. We evaluated the effect of sex on CHD as mediated by metabolomics or gene expression, adjusting for age and smoking status (never/ever) at the exam when the blood sample was collected. Of note, in our proposed multiple-mediator analysis framework, we include the selected omics mediators after variable selection in a multiple logistic regression model (equation (3)), so that the mediators can adjust for each other as potential confounders (VanderWeele et al., 2014). Here we rule out the possibility that the omics mediators could be colliders by imposing the temporal order among the exposure, mediators and the outcome. A collider is a variable that is caused by both the exposure and the outcome. Inappropriate adjustment of colliders in regression models can introduce bias (Holmberg and Andersen, 2022). Our proposed framework incorporates temporal ordering to ensure that putative mediators occur after the exposure and before the outcome, i.e., sex → transcriptome or metabolome → incident CHD. Therefore, the omics mediators cannot be colliders.

In addition, since the FHS employed a family-based study design, molecular phenotypes, such as gene expression and metabolites, can be correlated due to shared genetics and environmental components within a family. To account for familial correlation in gene expression and metabolite data prior to mediation analysis, we employed a linear mixed model (LMM) framework in which family structure was modeled as a random effect to remove correlations due to family relatedness (Cao et al., 2015). Specifically, we fit a LMM and specified the random effect covariance structure to be proportional to the family kinship matrix, which captures expected genetic relatedness among individuals. The model is defined as: *m*_*i*_ = *β*_0_ + δ_*i*_ + *ε*_*i*_, where *m*_*i*_ is a gene’s or metabolite’s level for individual *i*, δ_*i*_ is the family random effect modeled as 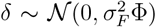, and *ε*_*i*_ ∼ 𝒩 (0, *σ*^2^) is the residual error. Here, Φ denotes the kinship matrix derived from pedigree information, representing the expected genetic correlation between individuals.

After fitting the model, we extracted the best linear unbiased prediction (BLUP) of the family random effect and subtracted it from each individual’s expression value to obtain an adjusted molecular mediator level: 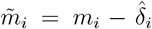 This transformation effectively removes family-related dependencies, yielding approximately independent samples suitable for downstream mediation analysis methods that assume independence (Cao et al., 2015).

We randomly split the data into 75% as the variable selection set and 25% as the estimation set. We used the first portion to select mediators and used the second portion to estimate the mediation effect. We also applied the product measure, difference measure and their proportion measures for comparison.

Metabolites have been shown to be associated with the risk of incident CHD (Wang et al., 2019)(Wang et al., 2023). However, the mediation effects of metabolites on the sex-CHD relationship have yet to be elucidated. Therefore, we applied MELODY to the plasma-based metabolomics data from individuals in the FHS Offspring Cohort. We evaluated the effect of sex on CHD risk as mediated by metabolites, adjusting for age and smoking status. As shown in Table 1, the total mediation effect of the metabolites was 0.014 (SE = 0.0007) by the *MED* measure. Because the values of *MED* measures depend on the total effect, we focused on more interpretable *SOS* measures in real data applications, which are standardized by the total effect and can be interpreted as the percentage of the total effect that the mediators explain. Based on the *SOS* measure, we estimated that 93% (SE = 5.0%) of the total effect of sex on incident CHD was mediated by metabolites. We identified five metabolites as mediators: dimethylglycine, carnitine, histidine, C48:0 triacylglycerol (TAG) and C16:1 sphingomyelins (SM). of note, it has been reported that histidine has protective effect on CHD risk (Yu et al., 2015), whereas dimethylglycine, carnitine, C48:0 TAG and C16:1 SM have been reported to be associated with incident CHD or its risk factors, such as obesity, fasting glucose, triglycerides and blood pressure (Ho et al., 2016; Wang et al., 2023). To our knowledge, this mediation analysis is the first to reveal how sex may influence the risk of CHD through these metabolites.

**Table 1.** Mediation effects of metabolomics (Metab) and gene expression (GE) on the sex-coronary heart disease (CHD) risk association by the proposed MELODY and product/difference measures. m: number of selected mediators. prop: proportion of the mediation effect relative to the total effect *c*.

| Mediator(s) | m | $R^2_{YX\vert Z}$ | $R^2_{YM\vert Z}$ | $R^2_{YMX\vert Z}$ | *MED* | *SOS* | *ab* | *ab(prop)* | $c - r$ | $c - r(\text{prop})$ |
| --- | --- | --- | --- | --- | --- | --- | --- | --- | --- | --- |
| *Multiple-mediator models* |  |  |  |  |  |  |  |  |  |  |
| Metab | 5 | 0.015 | 0.075 | 0.076 | 0.014 | 0.93 | -0.24 | 0.77 | -0.22 | 0.70 |
| GE | 10 | 0.015 | 0.040 | 0.051 | 0.0041 | 0.27 | -0.028 | 0.076 | -0.034 | 0.091 |
| *Single-mediator models for Metab* |  |  |  |  |  |  |  |  |  |  |
| dimethylglycine | 1 | 0.015 | 0.028 | 0.037 | 0.0068 | 0.46 | -0.082 | 0.26 | -0.073 | 0.23 |
| carnitine | 1 | 0.015 | 0.0046 | 0.017 | 0.0031 | 0.21 | -0.028 | 0.087 | -0.025 | 0.078 |
| histidine | 1 | 0.015 | 0.0052 | 0.020 | 0.00064 | 0.043 | -0.0090 | 0.029 | -0.0058 | 0.019 |
| C48:0 triacylglycerol | 1 | 0.015 | 0.0020 | 0.017 | 0.00047 | 0.031 | -0.0071 | 0.023 | -0.0044 | 0.014 |
| C16:1 sphingomyelins | 1 | 0.015 | 0.0094 | 0.019 | 0.0060 | 0.40 | -0.056 | 0.18 | -0.055 | 0.17 |

For the gene expression data, we identified 10 genes as mediators of the effect of sex on CHD risk, including HEMGN, FUCA2, TXK, SLC22A15, SLC4A10, IGJ, NRCAM, GNAO1, CEACAM8, and CD24. The total mediation effect of gene expression was 0.0041 (nonparametric Bootstrap-based SE = 0.0006) on CHD risk by the *MED* measure. We estimated that 27.1% (SE = 4%) of the total effect of sex on CHD risk was mediated by gene expressions based on the *SOS* measure. In addition, as shown in Table 1, the product and difference measures differed by around 19%, indicating inconsistency between the two measures. Between the gene expression and metabolites, the latter mediated more on the effect of sex on incident CHD (*SOS* = 0.27 and 0.93, respectively), suggesting that the metabolome is likely more pertinent to the CHD, a cardiometabolic disease (Wang et al., 2023).

## 5 Discussion

We have developed MELODY, a novel mediation analysis method, for high-dimensional mediators and binary outcomes to estimate the total mediation effect and identify mediators. MELODY critically extends the existing *R*^2^-based mediation analysis to binary outcomes in the high-dimensional setting. It performs well in mediation effect estimation and mediator selection, which we have shown via simulation studies and multiple real data applications. We would like to remind the readers that including U- or V-type non-mediators in estimating 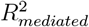 could result in substantial bias, as illustrated by Table S4, while our proposed mediator selection procedure can effectively mitigate the bias. Extending the *R*^2^ measure in linear regression to logistic regression is not straightforward owing to the nonunique definition of pseudo-*R*^2^ as we elaborated on. Based on our simulations, McFadden’s *R*^2^-based measure should be chosen for logistic regression models because the influence of disease prevalence on it is less than that on other pseudo-*R*^2^ measures.

Many questions of interest about mediation analysis for high-dimensional binary data remain to be addressed in the future. Mediation analysis of multiple types of high-dimensional omics mediators, such as DNA methylation in addition to gene expression and metabolomics, is biologically interesting yet analytically challenging due to the need for selecting mediators from multi-omics data with potentially complex dependence structures (Wang and Wei, 2020). Although we have proposed using the partial *R*^2^ measure for MELODY to control confounding in mediation analysis, models allowing for exposure-mediator interactions warrant further investigation in the future.

R functions that implement the proposed MELODY method are available on GitHub and will be available as an R package soon: https://github.com/inteomics/MELODY

## Acknowledgments

This work was supported by the National Institutes of Health (NIH) grants R01HL116720 and R21HL170213. P.W. was partially supported by NIH grants P50CA217674, P01CA296429 and Cancer Prevention and Research Institute of Texas (CPRIT) Grant RP230166. X.H. was partially supported by NIH grants R01CA272806, U54CA096300, U01CA152958 and P50CA100632 and the Dr. Mien-Chie Hung and Mrs. Kinglan Hung Endowed Professorship.

## Supplement to “MELODY: Mediation Analysis in Logistic Regression for High-Dimensional Mediators and a Binary Outcome”

The Supplementary Web materials contain the complete results of the simulation study discussed in the main text.

## References

Allison, P. D. (2014). Measures of fit for logistic regression. SAS Global Forum, Washington, DC.

Baron, R. M., Kenny, D. A. (1986). The Moderator-Mediator Variable Distinction in Social Psychological Research. Conceptual, Strategic, and Statistical Considerations. Journal of Personality and Social Psychology, 51(6).

Benjamini, Y., Hochberg, Y. (1995). Controlling the False Discovery Rate: A Practical and Powerful Approach to Multiple Testing. Journal of the Royal Statistical Society: Series B (Methodological), 57(1).

Cao, Y., Maxwell, T.J., Wei, P. (2015) A family-based joint test for mean and variance heterogeneity for quantitative traits. Annals of Human Genetics, 79(1):46–56.

Cashin, A.G, McAuley, J.H., VanderWeele, T.J., Lee, H. (2021). Understanding how health interventions or exposures produce their effects using mediation analysis. BMJ, 382:e071757.

Caubet, M., Samoilenko, M, Drouin, S., Sinnett, D., Krajinovic, M., Laverdi’ere, C., Marcil, V., Lefebvre, G.. (2023). Bayesian joint modeling for causal mediation analysis with a binary outcome and a binary mediator: Exploring the role of obesity in the association between cranial radiation therapy for childhood acute lymphoblastic leukemia treatment and the long-term risk of insulin resistance, Computational Statistics & Data Analysis, 10.1016/j.csda.2022.107586, 177, 107586.

Cavus, E., Karakas, M., Ojeda, F. M., Kontto, J., Veronesi, G., Ferrario, M. et al. (2019). Association of Circulating Metabolites With Risk of Coronary Heart Disease in a European Population: Results From the Biomarkers for Cardiovascular Risk Assessment in Europe (BiomarCaRE) Consortium. JAMA Cardiology, 4(12):1270–1279.

Chi, S., Flowers, C., Li, Z., Huang, X., Wei, P. (2024). MASH: mediation analysis of survival outcome and high-dimensional omics mediators with application to complex diseases. Annals of Applied Statistics, 18(2):1360–1377.

Clark-Boucher, D., Zhou, X., Du, J., Liu, Y., Needham, B.L., Smith, J.A., Mukherjee, B. (2023) Methods for mediation analysis with high-dimensional DNA methylation data: Possible choices and comparisons. PLoS Genet 19(11): e1011022.

Derkach, A., Pfeiffer, R. M., Chen, T. H., Sampson, J. N. (2019). High dimensional mediation analysis with latent variables. Biometrics, 75(3).

Doretti, M., Raggi, M., Stanghellini, E. (2022). Exact parametric causal mediation analysis for a binary outcome with a binary mediator. Statistical Methods and Applications, 31(1).

Fairchild, A. J., Mackinnon, D. P., Taborga, M. P., Taylor, A. B. (2009). R2 effect-size measures for mediation analysis. Behavior research methods, 41(2), 486–498.

Fan, J., Lv, J. (2008). Sure independence screening for ultrahigh dimensional feature space. Journal of the Royal Statistical Society. Series B: Statistical Methodology, 70(5).

Fang, R., Yang, H., Gao, Y., Cao, H., Goode, E.L., Cui, Y. (2021) Gene-based mediation analysis in epigenetic studies, Briefings in Bioinformatics, 22(3):bbaa113.

Fritz, M. S., MacKinnon, D. P. (2008). A graphical representation of the mediated effect. Behavior Research Methods, 40(1).

Gaynor, S. M., Schwartz, J., Lin, X. (2019). Mediation analysis for common binary outcomes. Statistics in Medicine, 38(4).

Ho, J. E., Larson, M. G., Ghorbani, A., Cheng, S., Chen, M. H., Keyes, M., Rhee, E. P., Clish, C. B., Vasan, R. S., Gerszten, R. E., Wang, T. J. (2016). Metabolomic profiles of body mass index in the Framingham Heart Study reveal distinct cardiometabolic phenotypes. PLoS ONE, 11(2), e0148361.

Holmberg, M.J., Andersen, L.W. (2022) Collider Bias. Journal of the American Medical Association, 327(13), 1282–1283.

Hughes, G., Choudhury, R. A., McRoberts, N. (2019). Summary measures of predictive power associated with logistic regression models of disease risk. Phytopathology, 109(5), 712–715.

Imai, K., Keele, L., Tingley, D. (2010). A general approach to causal mediation analysis. Psychological methods, 15(4), 309–334.

Kvalseth, T. O. (1985). Cautionary Note about R 2. The American Statistician, 39(4).

Lai, E. Y., Shih, S., Huang, Y. T., Wang, S. (2020). A mediation analysis for a nonrare dichotomous outcome with sequentially ordered multiple mediators. Statistics in Medicine, 39(10).

Lindenberger, U., Pötter, U. (1998). The Complex Nature of Unique and Shared Effects in Hierarchical Linear Regression: Implications for Developmental Psychology. Psychological Methods, 3(2).

Liu, Z., Shen, J., Barfield, R., Schwartz, J., Baccarelli, A. A., Lin, X. (2022). Large-Scale Hypothesis Testing for Causal Mediation Effects with Applications in Genome-wide Epigenetic Studies. Journal of the American Statistical Association, 117(537).

Loh, W. W., Moerkerke, B., Loeys, T., Vansteelandt, S. (2022). Nonlinear mediation analysis with high-dimensional mediators whose causal structure is unknown. Biometrics, 78(1).

MacKinnon, D.P. (2008) Introduction to Statistical Mediation Analysis. First Edition. New York: Taylor and Francis.

Menard, S. (2000). Coefficients of Determination for Multiple Logistic Regression Analysis (Vol. 54, Issue 1).

McFadden, D. (1973). Conditional logit analysis of qualitative choice behaviour. In P. Zarembka (Ed.),. In Frontiers of Econometrics. New York: Academic Press.

Nagelkerke, N. J. D. (1991). A note on a general definition of the coefficient of determination. In Biometrika (Vol. 78, Issue 3).

Royston, P. (2006). Explained variation for survival models. In Stata Journal (Vol. 6, Issue 1).

Nguyen, T. Q., Webb-Vargas, Y., Koning, I. M., Stuart, E. A. (2016). Causal Mediation Analysis With a Binary Outcome and Multiple Continuous or Ordinal Mediators: Simulations and Application to an Alcohol Intervention. Structural Equation Modeling, 23(3).

Pearl, J. (2001). Direct and indirect effects. In Proceedings of the Seventeenth Conference on Uncertainty and Artificial Intelligence, pp. 411–420, San Francisco. Morgan Kaufmann.

Rijnhart, J.J.M., Twisk, J.W.R., Eekhout, I. et al. (2019). Comparison of logisticregression based methods for simple mediation analysis with a dichotomous outcome variable. BMC Med Res Methodol 19, 19.

Robins, J. M., Greenland, S. (1992). Identifiability and exchangeability for direct and indirect effects. Epidemiology, 3(2).

Samoilenko, M., Lefebvre, G. (2021). Parametric-Regression-Based Causal Mediation Analysis of Binary Outcomes and Binary Mediators: Moving beyond the Rareness or Commonness of the Outcome. American Journal of Epidemiology, 190(9).

Schemper, M., Stare, J. (1996). Explained variation in survival analysis. Statistics in Medicine, 15(19).

Shi, B., Huang, X., Wei, P. (2022). Comparison of Effect Size Measures for Mediation Analysis of Survival Outcomes with Application to the Framingham Heart Study. 2205.03303.

Sohn, M., Li, H. (2019) Compositional mediation analysis for microbiome studies. Annals of Applied Statistics, 13:661–681.

Sohn, M.B., Lu, J., Li, H. (2022) A compositional mediation model for a binary outcome: Application to microbiome studies, Bioinformatics, 38(1):16–21.

Splansky, G.L., Corey, D., Yang, Q., Atwood, L.D., Cupples, L.A., Benjamin, E.J., et al. (2007) The Third Generation Cohort of the National Heart, Lung, and Blood Institute’s Framingham Heart Study: Design, Recruitment, and Initial Examination. American Journal of Epidemiology, 165(11):1328–1335.

Tjur, T. (2009). Coefficients of determination in logistic regression models A new proposal: The coefficient of discrimination. American Statistician, 63(4), 366–372.

Tsao, C.W., A.W. Aday, Z.I. Almarzooq, A. Alonso, A.Z. Beaton, M.S. Bittencourt, et al. (2022) Heart Disease and Stroke Statistics-2022 Update: A Report From the American Heart Association. Circulation, 145(8): p. e153–e639.

Valeri, L., and Vanderweele, T. J. (2013). Mediation analysis allowing for exposure-mediator interactions and causal interpretation: theoretical assumptions and implementation with SAS and SPSS macros. Psychological methods, 18(2), 137–150.

VanderWeele, T. J., Vansteelandt, S. (2010). Odds ratios for mediation analysis for a dichotomous outcome. American Journal of Epidemiology, 172(12).

VanderWeele, T. J., Vansteelandt, S. (2014). Mediation Analysis with Multiple Mediators. Epidemiologic methods, 2(1), 95–115.

VanderWeele, T. J. (2016). Mediation Analysis: A Practitioner’s Guide. In Annual Review of Public Health (Vol. 37).

Wang, K. (2018) Understanding Power Anomalies in Mediation Analysis. Psychometrika, 83(2):387–406.

Wang, W., Nelson, S., Albert, J. M. (2013). Estimation of causal mediation effects for a dichotomous outcome in multiple-mediator models using the mediation formula. Statistics in Medicine, 32(24).

Wang, Z., Wei, P. (2020) IMIX: a multivariate mixture model approach to assciation analysis through multi-omics data integration, Bioinformatics, 36 (22-23):5439–5447.

Wang, Z., Zhu, C., Nambi, V., Morrison, A.C., Folsom, A.R., Ballantyne, C.M., Boerwinkle, E., Yu, B. (2019). Metabolomic Pattern Predicts Incident Coronary Heart Disease. Arterioscler Thromb Vasc Biol., 39(7):1475–1482.

Wang, F., Tessier, A. J., Liang, L., Wittenbecher, C., Haslam, D. E., Fernández-Duval, G., Eliassen, H. A., Rexrode, K. M., Tobias, D. K., Li, J., Zeleznik, O., Grodstein, F., Martínez-González, M. A., Salas-Salvadó, J., Clish, C., Lee, K. H., Sun, Q., Stampfer, M. J., Hu, F. B., Guasch-Ferré, M. (2023). Plasma metabolomic profiles associated with mortality and longevity in a prospective analysis of 13,512 individuals. Nature Communications, 14(1), 5744.

Xu, Z., Li, C., Chi, S., Yang, T., Wei, P. (2024). Speeding up interval estimation for R2-based mediation effect of high-dimensional mediators via cross-fitting. Biostatistics. 2024 Oct 16:kxae037. doi: 10.1093/biostatistics/kxae037.

Xu, Z., Wei, P. (2024). A novel statistical framework for meta-analysis of total mediation effect with high-dimensional omics mediators in large-scale genomic consortia. PLOS Genetics. 20(11):e1011483. doi: 10.1371/journal.pgen.1011483.

Xue, F., Tang, X., Kim, G., Koenen, K. C., Martin, C. L., Galea, S., Wildman, D., Uddin, M., Qu, A. (2022). Heterogeneous Mediation Analysis on Epigenomic PTSD and Traumatic Stress in a Predominantly African American Cohort. Journal of the American Statistical Association, 117(540), 1669–1683.

Yang, T., Niu, J., Chen, H., Wei, P. (2021). Estimation of total mediation effect for high-dimensional omics mediators. BMC Bioinformatics, 22(1).

Yu, B., Li, A. H., Muzny, D., Veeraraghavan, N., de Vries, P. S., Bis, J. C., Musani, S. K., Alexander, D., Morrison, A. C., Franco, O. H., Uitterlinden, A., Hofman, A., Dehghan, A., Wilson, J. G., Psaty, B. M., Gibbs, R., Wei, P., Boerwinkle, E. (2015). Association of rare loss-of-function alleles in HAL, serum histidine levels and incident coronary heart disease. Circulation: Cardiovascular Genetics, 8(2), 351–355.

Zhang, C. H. (2010). Nearly unbiased variable selection under minimax concave penalty. Annals of Statistics, 38(2).

Zhang, J., Wei, Z., Chen, J. (2018) A distance-based approach for testing the mediation effect of the human microbiome, Bioinformatics, 34(11):1875–1883.

Zeng, P., Shao, Z., Zhou, X. (2021). Statistical methods for mediation analysis in the era of high-throughput genomics: Current successes and future challenges. In Computational and Structural Biotechnology Journal (Vol. 19).

Zhong, H., Prentice, R.L. (2008) Bias-reduced estimators and confidence intervals for odds ratios in genome-wide association studies. Biostatistics, 9(4):621–34.

