## Supplemental Materials for "MELODY: Mediation Analysis in Logistic Regression for High-Dimensional Mediators and a Binary Outcome"

### 1 Derivation of MED in Logistic models

$$R^2_{Y,MXZ} = 1 - \frac{\sum_{i=1}^{n}[y_i(\alpha_1 + rX + gZ + \sum_{j=1}^{p} b_j M_j) - log(1 + e^{\alpha_1 + rX + gZ + \sum_{j=1}^{p} b_j M_j})]}{\sum_{i=1}^{n}[y_i log(\bar{y}) + (1 - y_i) log(1 - \bar{y})]}, \quad (1)$$

$$R^2_{Y,MZ} = 1 - \frac{\sum_{i=1}^{n}[y_i(\alpha_2 + \sum_{j=1}^{p} d_j M_j + hZ) - log(1 + e^{\alpha_2 + \sum_{j=1}^{p} d_j M_j + hZ})]}{\sum_{i=1}^{n}[y_i log(\bar{y}) + (1 - y_i) log(1 - \bar{y})]}, \quad (2)$$

$$R^2_{Y,XZ} = 1 - \frac{\sum_{i=1}^{n}[y_i(\alpha_0 + cX + fZ) - log(1 + e^{\alpha_0 + cX + fZ})]}{\sum_{i=1}^{n}[y_i log(\bar{y}) + (1 - y_i) log(1 - \bar{y})]}, \quad (3)$$

$$R^2_{Y,Z} = 1 - \frac{\sum_{i=1}^{n}[y_i(\alpha_3 + kZ) - log(1 + e^{\alpha_3 + kZ})]}{\sum_{i=1}^{n}[y_i log(\bar{y}) + (1 - y_i) log(1 - \bar{y})]}, \quad (4)$$

$$\begin{aligned} MED &= R^2_{Y,M|Z} + R^2_{Y,X|Z} - R^2_{Y,MX|Z} = \frac{R^2_{Y,MZ} + R^2_{Y,XZ} - R^2_{Y,MXZ} - R^2_{Y,Z}}{1 - R^2_{Y,Z}} \\ &= \frac{\sum_{i=1}^{n}[y_i(\alpha_1 + rX + gZ + \sum_{j=1}^{p} b_j M_j + \alpha_3 + kZ - \alpha_2 - \sum_{j=1}^{p} d_j M_j - hZ - \alpha_0 - cX - fZ)]}{\sum_{i=1}^{n}[y_i(\alpha_3 + kZ) - log(1 + e^{\alpha_3 + kZ})]} \\ &\quad \frac{+ \sum_{i=1}^{n}[log\frac{(1+e^{\alpha_2 + \sum_{j=1}^{p} d_j M_j + hZ})(1+e^{\alpha_0 + cX + fZ})}{(1+e^{\alpha_1 + rX + gZ + \sum_{j=1}^{p} b_j M_j})(1+e^{\alpha_3 + kZ})}]}{} \end{aligned} \quad (5)$$

### 2 Comparison of pseudo-$R^2$ measures in Logistic regression for mediation analysis

This section elucidates the findings from our comparative analysis of three pseudo $R^2$ measures: McFadden's $R^2$, Nagelkerke's $R^2$, and Tjur's $R^2$, using comprehensive series of simulation studies. Simulation studies were structured in three main settings to evaluate the performance of McFadden's $R^2$, Nagelkerke's $R^2$, Tjur's $R^2$, product, and difference measures. For each setting, we present plots of $R^2$-based mediation effect measures that illustrate their performance and suitability of each $R^2$ measure in varying epidemiological scenarios. Firstly, as shown in Supplemental Figure 1, McFadden's $R^2$-based mediation measure exhibited stability and independence from disease prevalence, ranging from rare to common, compared with Nagelkerke's $R^2$, Tjur's $R^2$ and the difference measure, which is an appealing property for binary outcomes. Secondly, McFadden's $R^2$ measure consistently showed lower values across the board, indicating a more conservative estimate of variance explained, which aligns with its known properties in the literature (Supplemental Figures 1-3). Thirdly, as effected, all mediation effect measures under comparison increased with increasing mediation effect (parameter $b$ in equation (2)) and increasing number of mediators (Supplemental Figures 2 and 3, respectively). Finally, the difference between the product measure and the difference measure increased as the prevalence of disease increased up to 50%, which aligns with its rare disease assumption (Supplemental Figure 1). Based on the simulation study, we therefore advocate for the use of McFadden's $R^2$ in logistic models for more robust and stable mediation effect estimation.

### 3 Simulation studies

We performed a series of simulation studies to assess the performance of our proposed MELODY procedure. We use seven supplementary tables to present results of the seven simulation settings we conducted, where we evaluate our total mediation effect estimation performance in terms of the bias and variance. For the mediator selection methods, we not only evaluate their true positive rate (TPR) and false positive rate (FPR) but also examine their impacts on the bias and standard deviation (SD) of the mediation effects estimations. Settings in Table S1, S2, and S4-S6 represent the scenarios of high-dimensional settings with fixed sample size n = 2000 and number of potential mediators p = 1000, but varying

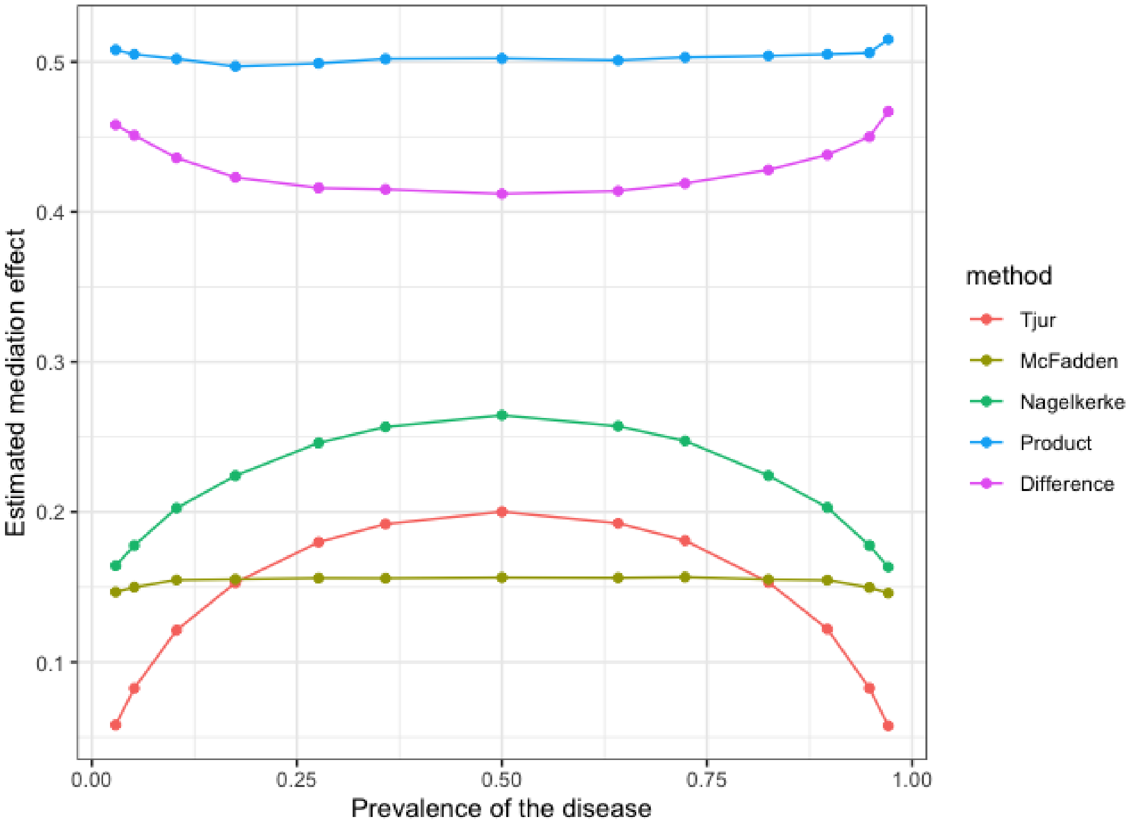


**Fig. 1** **Varying Prevalence of Disease.** By altering the prevalence from 0 to 1, we assessed the robustness of $R^2$ measures across different disease frequencies with the number of mediators fixed at 5. The resulting plot demonstrates each measure's stability or variability across varying disease prevalence.

the effect sizes of mediators, the prevalence of disease, the number of true mediators, the dimensions of data, and the confounding effect sizes. Table 3 showed the results of setting III where we introduce 10 U or V types non-mediators for each setting besides the noise variables to evaluate the effect of non-mediators U and V. In the settings VII presented in Table S7, we let the true mediators have different directions of mediation effects.

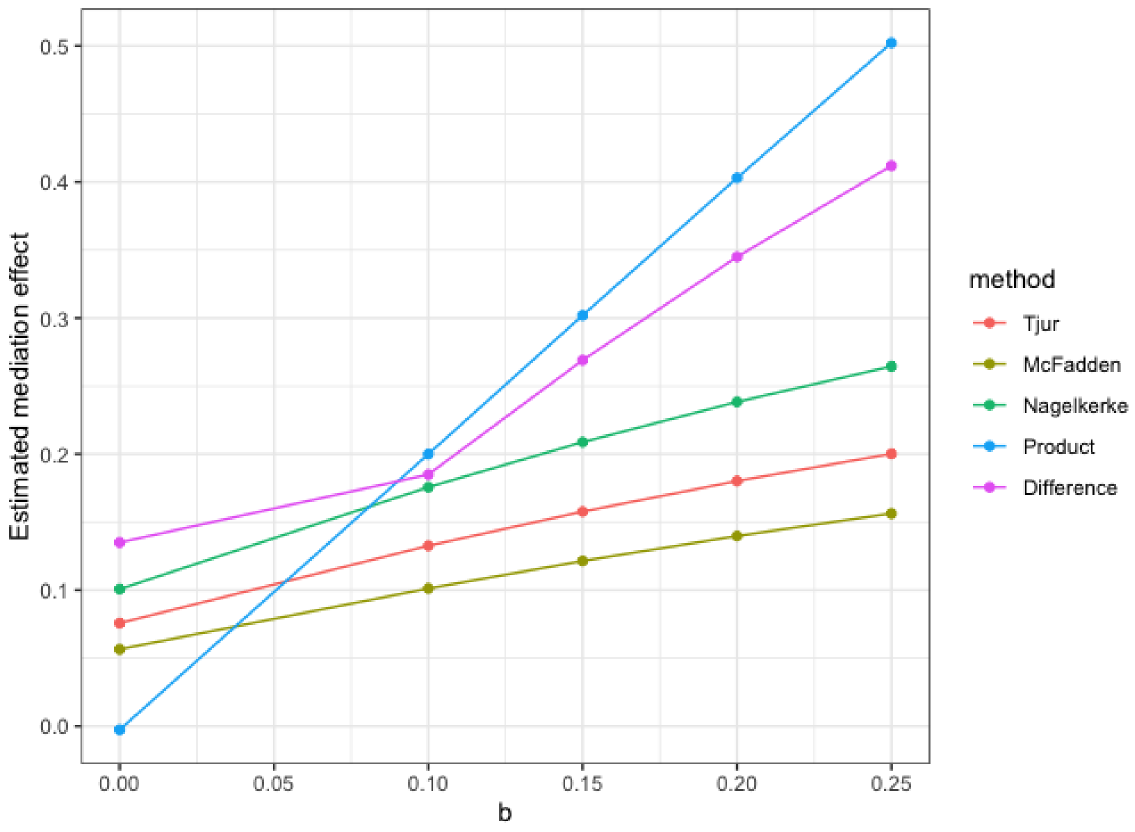


**Fig. 2 Varying Effect Sizes.** We examined how the $R^2$ measures respond to changes in effect sizes by adjusting the coefficient $b$ in equation (2). Plots depicting this relationship showcase the sensitivity of each $R^2$ measure to the strength of mediation effects.

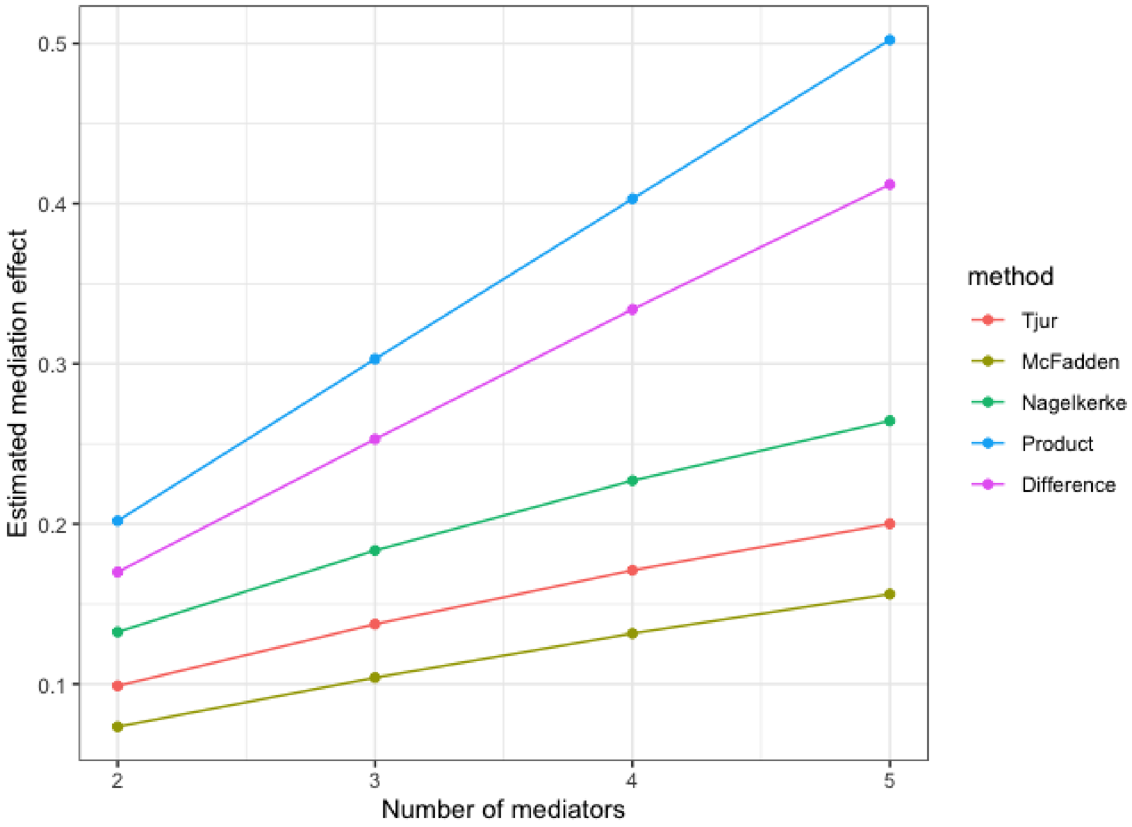


**Fig. 3 Varying the Number of Mediators** We explored the impact of the number of mediators on $R^2$ measures. The plots from this setting reveal the scalability of the $R^2$ measures and their behavior as the complexity of the model increases.

**Table 1** Bias and standard deviation under high-dimensional settings varying effect sizes

| Settings | Mediator Selection | | | Explained Variation | | Product | | Difference | |
|---|---|---|---|---|---|---|---|---|---|
| | Threshold | TPR | FPR | $MED$ | $SOS$ | $ab$ | Proportion | $c-r$ | Proportion |
| 1.1 | | | | | | | | | |
| $ab = 0.5$ | Null | 0.9839 | 0.0693 | -0.0053 (0.0246) | -0.0220 (0.0887) | 0.1157 (0.1976) | 0.0837 (0.1432) | -0.2852 (0.1965) | -0.1959 (0.1403) |
| | 0.2 | 0.9839 | 0.0044 | -0.0035 (0.0229) | -0.0139 (0.0764) | -0.0032 (0.1207) | 2e-04 (0.0935) | -0.0338 (0.0907) | -0.0216 (0.0713) |
| | 0.05 | 0.9839 | 0.0007 | -0.0035 (0.0228) | -0.0141 (0.0755) | -0.0085 (0.1179) | -0.0036 (0.0913) | -0.0248 (0.0870) | -0.0154 (0.0687) |
| 1.2 | | | | | | | | | |
| $ab = 0.4$ | Null | 0.9787 | 0.0809 | -0.0064 (0.0258) | -0.0281 (0.1049) | 0.1008 (0.1848) | 0.0771 (0.1402) | -0.3135 (0.1734) | -0.2282 (0.1294) |
| | 0.2 | 0.9783 | 0.0052 | -0.0042 (0.0241) | -0.0175 (0.0934) | -0.0019 (0.1163) | 9e-04 (0.0916) | -0.0346 (0.0905) | -0.0236 (0.0725) |
| | 0.05 | 0.9783 | 0.0008 | -0.0042 (0.024) | -0.0175 (0.0929) | -0.0069 (0.1133) | -0.0029 (0.0891) | -0.0241 (0.0886) | -0.0158 (0.071) |
| 1.3 | | | | | | | | | |
| $ab = 0.3$ | Null | 0.9238 | 0.0849 | -0.0133 (0.0300) | -0.0663 (0.1434) | 0.0437 (0.1727) | 0.0366 (0.1392) | -0.3420 (0.1711) | -0.2646 (0.1331) |
| | 0.2 | 0.9238 | 0.0052 | -0.0106 (0.0286) | -0.0523 (0.1335) | -0.0267 (0.1170) | -0.0188 (0.0958) | -0.0532 (0.0975) | -0.0399 (0.0804) |
| | 0.05 | 0.9238 | 0.0009 | -0.0107 (0.0285) | -0.0526 (0.1331) | -0.0306 (0.1148) | -0.0219 (0.0940) | -0.0439 (0.0960) | -0.0324 (0.0792) |
| 1.4 | | | | | | | | | |
| $ab = 0.2$ | Null | 0.8136 | 0.0870 | -0.0198 (0.029) | -0.1078 (0.1602) | -0.0021 (0.1504) | 9e-04 (0.1300) | -0.3395 (0.1614) | -0.2819 (0.1322) |
| | 0.2 | 0.8136 | 0.0045 | -0.0176 (0.0276) | -0.0963 (0.1476) | -0.0428 (0.1030) | -0.0340 (0.0895) | -0.0628 (0.0904) | -0.0511 (0.0786) |
| | 0.05 | 0.8136 | 0.0008 | -0.0175 (0.0275) | -0.0957 (0.1476) | -0.0438 (0.1017) | -0.0349 (0.0881) | -0.0538 (0.0899) | -0.0434 (0.0782) |
| 1.5 | | | | | | | | | |
| $ab = 0.1$ | Null | 0.8246 | 0.0885 | -0.0101 (0.0187) | -0.0641 (0.1198) | 0.0016 (0.0926) | 0.0024 (0.0858) | -0.3171 (0.125) | -0.2878 (0.1053) |
| | 0.2 | 0.8246 | 0.0048 | -0.0083 (0.0140) | -0.0531 (0.0847) | -0.0202 (0.0510) | -0.0180 (0.0469) | -0.0390 (0.0435) | -0.0352 (0.0403) |
| | 0.05 | 0.8246 | 0.0009 | -0.0083 (0.0138) | -0.0528 (0.0838) | -0.0208 (0.0502) | -0.0186 (0.0461) | -0.0299 (0.0411) | -0.0269 (0.0380) |

Setting I: Varying effect sizes (ab). We set the sample size $n = 2000$, number of true mediators $m = 5$, number of potential mediators $p = 1000$, $r = 1$, $\alpha_1 = 0$, we vary value of coefficients $a$ and $b$ in setting I s.t. $ab = 0.5, 0.4, 0.3, 0.2, 0.1$, respectively. We evaluate the mediator selection methods in tree thresholds: Null (SIS+MCP and not constraint FDR), 0.2 (SIS+MCP and constraint FDR <0.2), and 0.05 (SIS+MCP and constraint FDR <0.05).

**Table 2** Bias and standard deviation under high-dimensional settings varying the prevalence of the disease

| Settings | Mediator Selection | | | Explained Variation | | Product | | Difference | |
|---|---|---|---|---|---|---|---|---|---|
| | Threshold | TPR | FPR | $MED$ | $SOS$ | $ab$ | Proportion | $c-r$ | Proportion |
| 2.1 | | | | | | | | | |
| $pr = 0.029$ | No variable selection | 1 | 1 | -0.0028 (0.0072) | -0.0055 (0.0175) | 0.0628 (0.0383) | 0.0459 (0.0280) | -0.2484 (0.0263) | 1.1441 (0.0422) |
| | No FDR control | 1.0000 | 0.0697 | -0.0021 (0.0118) | -9e-04 (0.0354) | 0.0127 (0.0704) | 0.0118 (0.0503) | -0.1705 (0.0456) | 1.1451 (0.0723) |
| | FDR<0.2 | 1.0000 | 0.0013 | -0.0021 (0.0118) | -0.0011 (0.0344) | -0.0027 (0.0662) | 0.0013 (0.0477) | -0.1513 (0.0423) | 1.1451 (0.0723) |
| | FDR<0.05 | 1.0000 | 0.0003 | -0.0021 (0.0118) | -0.0010 (0.0343) | -0.0028 (0.0661) | 0.0013 (0.0476) | -0.1510 (0.0422) | 1.1451 (0.0723) |
| 2.2 | | | | | | | | | |
| $pr = 0.052$ | No variable selection | 1 | 1 | -0.0011 (0.0057) | -0.0066 (0.0215) | 0.1080 (0.0435) | 0.0741 (0.0317) | -0.2843 (0.0334) | 1.1460 (0.0384) |
| | No FDR control | 0.9996 | 0.0774 | -4e-04 (0.0129) | 2e-04 (0.036) | 0.0299 (0.0718) | 0.0218 (0.0526) | -0.1742 (0.0467) | 1.1442 (0.0833) |
| | FDR<0.2 | 0.9996 | 0.0017 | -3e-04 (0.0127) | 6e-04 (0.0353) | 0.0062 (0.0651) | 0.0055 (0.0484) | -0.1427 (0.0419) | 1.1442 (0.0833) |
| | FDR<0.05 | 0.9996 | 0.0003 | -3e-04 (0.0127) | 7e-04 (0.0353) | 0.0059 (0.0649) | 0.0053 (0.0482) | -0.1422 (0.0418) | 1.1442 (0.0833) |
| 2.3 | | | | | | | | | |
| $pr = 0.175$ | No variable selection | 1 | 1 | -0.0194 (0.0315) | -0.0047 (0.0088) | 0.2836 (0.0879) | 0.1989 (0.0632) | -0.5283 (0.0677) | 1.1338 (0.0542) |
| | No FDR control | 0.9992 | 0.0816 | -7e-04 (0.0493) | -1e-04 (0.0158) | 0.0610 (0.0958) | 0.0428 (0.071) | -0.2049 (0.0647) | 1.1406 (0.1035) |
| | FDR<0.2 | 0.9992 | 0.0037 | -0.0011 (0.0455) | -2e-04 (0.0152) | 0.0066 (0.0805) | 0.0050 (0.0610) | -0.1351 (0.0493) | 1.1406 (0.1035) |
| | FDR<0.05 | 0.9744 | 0.0003 | -0.0137 (0.0680) | -0.0029 (0.0187) | -0.0092 (0.0916) | -0.0060 (0.0678) | -0.1410 (0.0558) | 1.1375 (0.1217) |
| 2.4 | | | | | | | | | |
| $pr = 0.276$ | No variable selection | 1 | 1 | -0.0125 (0.0109) | -0.0558 (0.0466) | 0.6294 (0.1879) | 0.4428 (0.1269) | -1.0090 (0.1908) | 1.1245 (0.0630) |
| | No FDR control | 0.9984 | 0.0837 | 0.0011 (0.0190) | -0.0022 (0.0576) | 0.0970 (0.1264) | 0.0661 (0.0917) | -0.2364 (0.0835) | 1.1389 (0.1234) |
| | FDR<0.2 | 0.9984 | 0.0042 | 0.0011 (0.0179) | -0.0019 (0.0493) | 0.0133 (0.0920) | 0.0076 (0.0680) | -0.1322 (0.0519) | 1.1389 (0.1234) |
| | FDR<0.05 | 0.9984 | 0.0008 | 0.0011 (0.0178) | -0.0021 (0.0492) | 0.0106 (0.0915) | 0.0057 (0.0677) | -0.1290 (0.0516) | 1.1389 (0.1234) |
| 2.5 | | | | | | | | | |
| $pr = 0.358$ | No variable selection | 1 | 1 | -0.1005 (0.0570) | -0.4449 (0.2639) | 1.6895 (4.1742) | 1.2076 (2.8903) | -3.1693 (7.0299) | 0.4415 (0.6987) |
| | No FDR control | 0.9980 | 0.0701 | -0.0027 (0.0191) | -0.0097 (0.0634) | 0.0788 (0.1292) | 0.0570 (0.0959) | -0.2457 (0.0833) | 1.1303 (0.1350) |
| | FDR<0.2 | 0.9980 | 0.0053 | -0.0017 (0.0185) | -0.0056 (0.0548) | 0.0061 (0.0988) | 0.0057 (0.0744) | -0.1384 (0.0565) | 1.1303 (0.1350) |
| | FDR<0.05 | 0.9980 | 0.0008 | -0.0017 (0.0185) | -0.0056 (0.0548) | 0.0017 (0.0975) | 0.0026 (0.0734) | -0.1330 (0.0562) | 1.1303 (0.1350) |

Setting II: Varying prevalence of the disease (pr). We set the sample size $n = 2000$, number of true mediators $m = 5$, number of potential mediators $p = 1000$, $r = 1$. For each true mediator, we set the $a = 0.4$ and $b = 0.25$. We vary value of coefficients $\alpha_1 = log(\frac{1}{100})$, $log(\frac{1}{9})$, $log(\frac{3}{7})$, 0, $log(\frac{7}{3})$, $log(9)$ s.t. $pr = 0.029$, 0.052, 0.175, 0.276, 0.358, respectively. We evaluate the mediator selection methods in four threshold levels: No variable selection, No FDR control (SIS+MCP only), FDR <0.2 (SIS+MCP and constraint FDR <0.2), and FDR <0.05 (SIS+MCP and constraint FDR <0.05).

**Table 3** Bias and standard deviation under high-dimensional settings with U or V type non-mediators exist

| Settings | Mediator Selection | | | Explained Variation | | Product | | Difference | |
|---|---|---|---|---|---|---|---|---|---|
| | Threshold | TPR | FPR | *MED* | *SOS* | *ab* | Proportion | $c - r$ | Proportion |
| 3.1 | | | | | | | | | |
| U type non-mediators | Null | 0.9958 | 0.0417 | -0.0261 (0.0211) | -0.0392 (0.0831) | 0.0924 (0.1476) | 0.1253 (0.1173) | -0.3224 (0.1095) | 0.9644 (0.1407) |
| | 0.2 | 0.9958 | 0.0046 | -0.0179 (0.0183) | 0.0067 (0.0652) | -0.0626 (0.097) | 9e-04 (0.0845) | -0.1121 (0.0606) | 0.9644 (0.1407) |
| | 0.05 | 0.9958 | 0.0009 | -0.0175 (0.0184) | 0.0085 (0.0650) | -0.0720 (0.0945) | -0.0068 (0.0820) | -0.1005 (0.0583) | 0.9644 (0.1407) |
| 3.2 | | | | | | | | | |
| V type non-mediators | Null | 0.9966 | 0.0291 | 0.0012 (0.0202) | 0.0035 (0.0705) | 0.0739 (0.1295) | 0.0544 (0.1043) | -0.1887 (0.0792) | 1.1123 (0.1521) |
| | 0.2 | 0.9966 | 0.0054 | 0.0019 (0.0194) | 0.0067 (0.0662) | 0.0297 (0.1099) | 0.0219 (0.0893) | -0.1312 (0.0625) | 1.1123 (0.1521) |
| | 0.05 | 0.9966 | 0.0021 | 0.0019 (0.0193) | 0.0069 (0.0656) | 0.0232 (0.1068) | 0.0172 (0.0874) | -0.1244 (0.0602) | 1.1123 (0.1521) |

Setting III: U or V types non-mediators exist. We set the sample size $n = 2000$, number of true mediators $m = 5$, number of potential mediators $p = 1000$, $r = 1$, and $\alpha_1 = 0$. For each true mediator, we set the $a = 0.2$ and $b = 0.3$. Except for noise variables, we introduce 10 U or V types non-mediators for each setting. For U type non-mediators, we set $a = 0$, $b = 0.3$. For V type non-mediators, we set $a = 0.2$, $b = 0$. We evaluate the mediator selection methods in tree thresholds: Null (SIS+MCP and not constraint FDR), 0.2 (SIS+MCP and constraint FDR <0.2), and 0.05 (SIS+MCP and constraint FDR <0.05).

**Table 4** Bias and standard deviation under high-dimensional settings varying the number of true mediators

| Settings | Mediator Selection | | | Explained Variation | | Product | | Difference | |
|---|---|---|---|---|---|---|---|---|---|
| | Threshold | TPR | FPR | *MED* | *SOS* | *ab* | Proportion | $c - r$ | Proportion |
| 4.1 | | | | | | | | | |
| $m = 5$ | Null | 0.9840 | 0.0694 | -0.0053 (0.0246) | -0.0220 (0.0887) | 0.1157 (0.1976) | 0.0837 (0.1432) | -0.2852 (0.1965) | -0.1959 (0.1403) |
| | 0.2 | 0.9840 | 0.0044 | -0.0035 (0.0229) | -0.0139 (0.0764) | -0.0032 (0.1207) | 2e-04 (0.0935) | -0.0338 (0.0907) | -0.0216 (0.0713) |
| | 0.05 | 0.9840 | 0.0007 | -0.0035 (0.0228) | -0.0141 (0.0755) | -0.0085 (0.1179) | -0.0036 (0.0913) | -0.0248 (0.0870) | -0.0154 (0.0687) |
| 4.2 | | | | | | | | | |
| $m = 20$ | Null | 0.9830 | 0.0329 | -0.0030 (0.0242) | -0.0144 (0.0581) | 0.1783 (0.2179) | 0.1070 (0.1424) | -0.2096 (0.1448) | -0.1241 (0.0987) |
| | 0.2 | 0.9830 | 0.0090 | -0.0024 (0.0238) | -0.0121 (0.0555) | 0.0777 (0.1820) | 0.0463 (0.1218) | -0.3188 (0.0803) | 1.2736 (0.1707) |
| | 0.05 | 0.9830 | 0.0023 | -0.0024 (0.0237) | -0.0119 (0.0546) | 0.0525 (0.1695) | 0.0311 (0.1143) | -0.3047 (0.0761) | 1.2736 (0.1707) |
| 4.3 | | | | | | | | | |
| $m = 50$ | Null | 0.9602 | 0.0132 | -0.0062 (0.0234) | -0.0312 (0.0699) | 0.2685 (0.2600) | 0.1696 (0.1725) | -0.3964 (0.1356) | 1.2090 (0.1612) |
| | 0.2 | 0.9556 | 0.0066 | -0.0064 (0.0231) | -0.0317 (0.0689) | 0.2092 (0.2438) | 0.1315 (0.1642) | -0.3685 (0.1244) | 1.2090 (0.1612) |
| | 0.05 | 0.9317 | 0.0026 | -0.0092 (0.0230) | -0.0433 (0.0693) | 0.1203 (0.2289) | 0.0743 (0.1573) | -0.3582 (0.1145) | 1.2090 (0.1612) |

Setting IV: Varying the number of true mediators. We set the sample size $n = 2000$, number of potential mediators $p = 1000$, $r = 1$, and $\alpha_1 = 0$. For each true mediator, we set the $a = 0.4$ and $b = 0.25$. We vary number of true mediators $m$ = 5, 20, 50. We evaluate the mediator selection methods in tree thresholds: Null (SIS+MCP and not constraint FDR), 0.2 (SIS+MCP and constraint FDR <0.2), and 0.05 (SIS+MCP and constraint FDR <0.05).

**Table 5** Bias and standard deviation under high-dimensional settings varying the dimension of data

| Settings | Mediator Selection | | | Explained Variation | | Product | | Difference | |
|---|---|---|---|---|---|---|---|---|---|
| | Threshold | TPR | FPR | $MED$ | $SOS$ | $ab$ | Proportion | $c-r$ | Proportion |
| 5.1 | | | | | | | | | |
| $p$ = 1000 | Null | 0.9839 | 0.0693 | -0.0053 (0.0246) | -0.0220 (0.0887) | 0.1157 (0.1976) | 0.0837 (0.1432) | -0.2852 (0.1965) | -0.1959 (0.1403) |
| | 0.2 | 0.9839 | 0.0044 | -0.0035 (0.0229) | -0.0139 (0.0764) | -0.0032 (0.1207) | 2e-04 (0.0935) | -0.0338 (0.0907) | -0.0216 (0.0713) |
| | 0.05 | 0.9839 | 0.0007 | -0.0035 (0.0228) | -0.0141 (0.0755) | -0.0085 (0.1179) | -0.0036 (0.0913) | -0.0248 (0.0870) | -0.0154 (0.0687) |
| 5.2 | | | | | | | | | |
| $p$ = 2000 | Null | 0.9976 | 0.0428 | -0.0042 (0.0235) | -0.0170 (0.0834) | 0.1574 (0.1958) | 0.1132 (0.1433) | -0.3477 (0.1958) | -0.2394 (0.1384) |
| | 0.2 | 0.9976 | 0.0057 | -0.0020 (0.0212) | -0.0076 (0.0658) | 0.0136 (0.1209) | 0.0120 (0.0938) | -0.0476 (0.0936) | -0.0313 (0.0735) |
| | 0.05 | 0.9976 | 0.0008 | -0.0019 (0.0210) | -0.0071 (0.0637) | 0.0012 (0.1139) | 0.0031 (0.0883) | -0.0212 (0.0838) | -0.0129 (0.0668) |
| 5.3 | | | | | | | | | |
| $p$ = 5000 | Null | 0.9960 | 0.0176 | -0.0037 (0.0234) | -0.0153 (0.0797) | 0.1696 (0.1910) | 0.1216 (0.1404) | -0.3590 (0.1836) | -0.2475 (0.1312) |
| | 0.2 | 0.9960 | 0.0046 | -0.0021 (0.0215) | -0.0078 (0.0682) | 0.0305 (0.1305) | 0.0239 (0.1005) | -0.0807 (0.1051) | -0.0541 (0.082) |
| | 0.05 | 0.9960 | 0.0007 | -0.0018 (0.0208) | -0.0067 (0.0635) | 0.0043 (0.1142) | 0.0054 (0.0890) | -0.0253 (0.0840) | -0.0157 (0.0673) |
| 5.4 | | | | | | | | | |
| $p$ = 10000 | Null | 0.9944 | 0.0089 | -0.0149 (0.0817) | -0.0036 (0.0241) | 0.1845 (0.1996) | 0.1322 (0.1465) | -0.3619 (0.1863) | -0.2500 (0.1324) |
| | 0.2 | 0.9944 | 0.0032 | -0.0098 (0.0682) | -0.0025 (0.0218) | 0.0449 (0.1344) | 0.0341 (0.1029) | -0.1116 (0.1120) | -0.0759 (0.0857) |
| | 0.05 | 0.9944 | 0.0006 | -0.0079 (0.0637) | -0.0021 (0.0210) | 0.0061 (0.1153) | 0.0066 (0.0892) | -0.0334 (0.0862) | -0.0214 (0.0681) |

Setting V: Varying the dimension of data (n/p). We set the sample size $n$ = 2000, number of true mediators $m$ = 5, $r$ = 1, and $\alpha_1 = 0$. For each true mediator, we set the $a$ = 0.4 and $b$ = 0.25. We vary number of potential mediators $p$ = 1000, 2000, 5000, and 10000. We evaluate the mediator selection methods in tree thresholds: Null (SIS+MCP and not constraint FDR), 0.2 (SIS+MCP and constraint FDR <0.2), and 0.05 (SIS+MCP and constraint FDR <0.05).

**Table 6** Bias and standard deviation under high-dimensional settings varying the confounding effects

| Settings | Mediator Selection | | | Explained Variation | | Product | | Difference | |
|---|---|---|---|---|---|---|---|---|---|
| | Threshold | TPR | FPR | *MED* | *SOS* | *ab* | Proportion | $c-r$ | Proportion |
| 6.1 | | | | | | | | | |
| $g = 0.1$ | Null | 0.9812 | 0.0730 | -0.0172 (0.0826) | -0.0050 (0.0218) | 0.1213 (0.1743) | 0.0901 (0.1352) | -0.3074 (0.1294) | 1.1193 (0.1472) |
| | 0.2 | 0.9812 | 0.0046 | -0.0127 (0.073) | -0.004 (0.0211) | 0.0013 (0.1117) | 0.0039 (0.0913) | -0.1317 (0.0715) | 1.1193 (0.1472) |
| | 0.05 | 0.9812 | 0.0008 | -0.0125 (0.0721) | -0.0039 (0.021) | -0.0029 (0.1098) | 9e-04 (0.0896) | -0.1251 (0.0693) | 1.1193 (0.1472) |
| 6.2 | | | | | | | | | |
| $g = 0.5$ | Null | 0.9924 | 0.0503 | -0.0129 (0.0709) | -0.0043 (0.0222) | 0.0703 (0.1384) | 0.0533 (0.1049) | -0.2359 (0.0922) | 1.1131 (0.1454) |
| | 0.2 | 0.9924 | 0.0054 | -0.0080 (0.0636) | -0.0032 (0.0213) | 0.0056 (0.1073) | 0.0068 (0.0834) | -0.1317 (0.0658) | 1.1131 (0.1454) |
| | 0.05 | 0.9924 | 0.0010 | -0.0074 (0.0628) | -0.0031 (0.0211) | 0.0010 (0.1045) | 0.0035 (0.0818) | -0.1233 (0.0632) | 1.1131 (0.1454) |
| 6.3 | | | | | | | | | |
| $g = 0.75$ | Null | 0.9872 | 0.0481 | -0.0191 (0.0813) | -0.0057 (0.0233) | 0.0579 (0.1481) | 0.0447 (0.1119) | -0.2401 (0.0985) | 1.1099 (0.1473) |
| | 0.2 | 0.9872 | 0.0047 | -0.0125 (0.0737) | -0.0043 (0.0222) | -2e-04 (0.1178) | 0.0029 (0.0905) | -0.1349 (0.0719) | 1.1099 (0.1473) |
| | 0.05 | 0.9872 | 0.0009 | -0.0125 (0.0731) | -0.0043 (0.022) | -0.0051 (0.1152) | -6e-04 (0.0890) | -0.1279 (0.0694) | 1.1099 (0.1473) |
| 6.4 | | | | | | | | | |
| $g = 1$ | Null | 0.9728 | 0.0449 | -0.0063 (0.0245) | -0.0197 (0.0870) | 0.0580 (0.1533) | 0.0460 (0.1163) | -0.2338 (0.1024) | 1.1069 (0.1510) |
| | 0.2 | 0.9724 | 0.0040 | -0.0056 (0.0243) | -0.0167 (0.0828) | -0.0042 (0.1227) | 0.0011 (0.0944) | -0.1366 (0.0739) | 1.1040 (0.1634) |
| | 0.05 | 0.9724 | 0.0008 | -0.0057 (0.0241) | -0.0169 (0.0821) | -0.0087 (0.1205) | -0.0022 (0.0928) | -0.1307 (0.0721) | 1.1040 (0.1634) |

Setting VI: Presence of confounding variables. We set the sample size $n = 2000$, number of true mediators $m = 5$, number of potential mediators $p = 1000$, $r = 1$, and $\alpha_1 = 0$. For each true mediator, we set the $a = 0.2$ and $b = 0.3$. For covariates $Z$, we set the parameters $l = 0.1$, and $g = 0.1$, 0.5, 0.75, and 1. We evaluate the mediator selection methods in tree thresholds: Null (SIS+MCP and not constraint FDR), 0.2 (SIS+MCP and constraint FDR <0.2), and 0.05 (SIS+MCP and constraint FDR <0.05).

**Table 7** Bias and standard deviation under high-dimensional settings with conflict direction effects

| Settings | Mediator Selection | | | Explained Variation | | Product | | Difference | |
|---|---|---|---|---|---|---|---|---|---|
| | Threshold | TPR | FPR | *MED* | *SOS* | *ab* | Proportion | $c-r$ | Proportion |
| 7.1 | | | | | | | | | |
| Same directions | Null | 0.9787 | 0.0809 | -0.0064 (0.0258) | -0.0281 (0.1049) | 0.1008 (0.1848) | 0.0771 (0.1402) | -0.3135 (0.1734) | -0.2282 (0.1294) |
| | 0.2 | 0.9783 | 0.0052 | -0.0042 (0.0241) | -0.0175 (0.0934) | -0.0019 (0.1163) | 9e-04 (0.0916) | -0.0346 (0.0905) | -0.0236 (0.0725) |
| | 0.05 | 0.9783 | 0.0008 | -0.0042 (0.0240) | -0.0175 (0.0929) | -0.0069 (0.1133) | -0.0029 (0.0891) | -0.0241 (0.0886) | -0.0158 (0.0710) |
| 7.2 | | | | | | | | | |
| Conflict directions | Null | 0.9860 | 0.0478 | -0.0016 (0.0214) | -0.0094 (0.0868) | 0.0597 (0.1528) | 0.0499 (0.1278) | -0.1635 (0.1269) | 1.0417 (0.1340) |
| | 0.2 | 0.9860 | 0.0045 | -9e-04 (0.0205) | -0.0055 (0.0792) | 0.0055 (0.1201) | 0.0055 (0.1023) | -0.0503 (0.0928) | 1.0417 (0.1340) |
| | 0.05 | 0.9860 | 0.0009 | -8e-04 (0.0205) | -0.0051 (0.0786) | 0.0031 (0.1181) | 0.0034 (0.1004) | -0.0422 (0.0903) | 1.0417 (0.1340) |

Setting VII: Conflict direction effects. We set the sample size n = 2000, number of true mediators m = 5, number of potential mediators p = 1000, r = 1, 1 = 0, we set the a = 0.4 for each true mediator. To study the scenario when mediators have different directions of effect, we set b = (0.5, 0.5, -0.25, -0.25, -0.25). We evaluate the mediator selection methods in three thresholds: Null (SIS+MCP and not constraint FDR), 0.2 (SIS+MCP and constraint FDR <0.2), and 0.05 (SIS+MCP and constraint FDR <0.05).

**Table 8** Bias and standard deviation under high-dimensional settings with correlated mediators

| Settings | Mediator Selection | | Explained Variation | | Product | | Difference | |
|---|---|---|---|---|---|---|---|---|
| | TPR | FPR | $MED$ | $SOS$ | $ab$ | Proportion | $c-r$ | Proportion |
| 8.1 | | | | | | | | |
| Random correlated | 0.9948 | 0.0062 | -5e-04<br>(0.0219) | -0.0097<br>(0.0626) | 0.0045<br>(0.1144) | 0.0013<br>(0.0837) | -1.035<br>(0.0855) | -0.1465<br>(0.0652) |
| Fixed correlated | 0.9932 | 0.0056 | -0.0023<br>(0.0185) | -0.0078<br>(0.0659) | 0.0051<br>(0.0954) | 0.0062<br>(0.0774) | -1.0315<br>(0.0636) | -0.1144<br>(0.0559) |
| 8.2 | | | | | | | | |
| Random correlated | 0.9974 | 0.0047 | -0.0186<br>(0.0178) | 0.0101<br>(0.0715) | -0.0633<br>(0.0937) | 0.0046<br>(0.0906) | -0.0513<br>(0.0673) | -0.1064<br>(0.0663) |
| Fixed correlated | 0.9664 | 0.0020 | -0.0721<br>(0.0160) | 0.0982<br>(0.1177) | 0.2448<br>(0.1455) | 0.6762<br>(0.1997) | -0.1689<br>(0.1274) | -0.0978<br>(0.1787) |

Setting VIII: Correlated mediators. In setting 8.1, we add correlation structures for mediators on setting 1; In setting 8.2, we add correlation structures for mediators on setting 3.1, where U type non-mediators exists.
